# A distinct cardiopharyngeal mesoderm genetic hierarchy establishes antero-posterior patterning of esophagus striated muscle

**DOI:** 10.1101/600841

**Authors:** Glenda Comai, Églantine Heude, Sebastien Mella, Sylvain Paisant, Francesca Pala, Mirialys Gallardo, Gabrielle Kardon, Swetha Gopalakrishnan, Shahragim Tajbakhsh

## Abstract

In most vertebrates, the upper digestive tract is composed of muscularised jaws linked to the esophagus that permit food uptake and swallowing. Masticatory and esophagus striated muscles (ESM) share a common cardiopharyngeal mesoderm (CPM) origin, however ESM are unusual among striated muscles as they are established in the absence of a primary skeletal muscle scaffold. Using mouse chimeras, we show that the transcription factors *Tbx1* and *Isl1* are required cell-autonomously for myogenic specification of ESM progenitors. Further, genetic loss-of-function and pharmacological studies point to Met/HGF signalling for antero-posterior migration of esophagus muscle progenitors, where HGF ligand is expressed in adjacent smooth muscle cells. These observations highlight the functional relevance of a smooth and striated muscle progenitor dialogue for ESM patterning. Our findings establish a *Tbx1-Isl1-Met* genetic hierarchy that uniquely regulate esophagus myogenesis and identify distinct genetic signatures that can be used as a framework to interpret pathologies arising within CPM derivatives.

## INTRODUCTION

Evolution of vertebrates has been marked by the emergence of muscularised jaws that transitioned them from filter feeders to active predators (Northcutt, 2005). Considerable diversity in developmental origins and regulation of skeletal muscles point to important functional differences that remain unexplored. Muscles of the trunk originate from the segmented somites, whereas head muscles arise independently, from the non-segmented cranial paraxial mesoderm located anterior to the somites (review Diogo & Sambasivan). The specification of head and trunk muscles involves divergent genetic regulatory networks, to activate bHLH myogenic regulatory factors (MRFs) Myf5, Mrf4, Myod and Myogenin that play crucial roles in governing striated muscle cell fate and differentiation (Comai et al., 2014; Kassar-Duchossoy et al., 2005; Rudnicki et al., 1993).

While somitic myogenic progenitors are regulated primarily by the paired/homeodomain genes *Pax3* and *Pax7* and *Myf5* that act genetically upstream of *Myod* (Kassar-Duchossoy et al., 2005; Relaix et al., 2005; Tajbakhsh et al., 1997), cardiopharyngeal mesoderm (CPM) progenitors, that colonize pharyngeal arches and form craniofacial and neck muscles, are regulated by a Pax3-independent regulatory network (Diogo et al., 2015; Sambasivan et al., 2011). CPM progenitors specified by *Tbx1* and *Isl1* genes are bipotent as they form branchiomeric subsets of head/neck muscles as well as the second heart field (Diogo et al., 2015; Kelly et al., 2004; Lescroart et al., 2015; Sambasivan et al., 2009). *Tbx1* acts together with *Myf5* to assure myogenic fate (Harel et al., 2009; Kelly et al., 2004; Nathan et al., 2008; Sambasivan et al., 2009). In Tbx1-null embryos, the first pharyngeal arch is hypoplastic and posterior pharyngeal arches do not form, resulting in variably penetrant defects of masticatory muscles and absence of muscles derived from more posterior arches including those of the larynx and esophagus (Gopalakrishnan et al., 2015; Heude et al., 2018; Kelly et al., 2004; Lescroart et al., 2015). *Tbx1* exerts cell-autonomous and non-autonomous roles as conditional deletion of *Tbx1* in CPM and pharyngeal endoderm phenocopies the pharyngeal arch and cardiac outflow tract phenotype of the null mutant (Arnold et al., 2006; Kelly et al., 2004; Zhang et al., 2006). On the other hand, the functional role of *Isl1* in CPM specification remains unknown due to early embryonic lethality of Isl1-null mutants (by E10.5) that exhibit cardiac deficiencies (Cai et al., 2003; Harel et al., 2009; Nathan et al., 2008). Thus, due to the severe phenotypes observed in the mouse, the epistatic relationship between *Tbx1* and *Isl1* and their cell-autonomous roles during ESM formation remain unclear.

Recent studies by us and others showed that CPM progenitors generate diverse myogenic subpopulations at the transition zone between head and trunk (Diogo et al., 2015; Gopalakrishnan et al., 2015; Heude et al., 2018; Schubert et al., 2018; Tabler et al., 2017). Whether CPM muscle derivatives form a homogeneous group specified by a unique gene regulatory network is unknown. Esophagus striated muscles (ESM) arise from CPM and they exhibit several features that are distinct from other striated muscles in the organism. Notably, ESM formation initiates in the foetus, thus embryonic myogenesis which generates primary myofibres that act as scaffolds for secondary (foetal) myofibres does not take place. As the esophagus is the only site identified to date that undergoes this unusual patterning, this raises the issue of what cell type (s) pattern ESM.

The esophageal muscularis externa comprises a variable amount of striated muscle (Krauss et al., 2016),which has a distinct developmental origin than the smooth muscle layer (Gopalakrishnan et al., 2015; Rishniw et al., 2003; Zhao and Dhoot, 2000). Interestingly, birds and reptiles lack the striated muscle component in the esophagus suggesting that mastication and processing of bolus through the digestive tract might have common evolutionary links (Gopalakrishnan et al., 2015). In ESM-containing species, postnatal maturation of the esophageal musculature involves proximo-distal replacement of smooth muscle by as yet elusive mechanisms (Krauss et al., 2016). Although smooth muscle and other mesenchymal cells are in close proximity to ESM progenitors as they undergo lineage commitment and differentiation, how the latter are patterned in the absence of primary myofibres remains unknown. It has been proposed that smooth muscle may provide a scaffold for laying down ESM myofibers, however it is unclear to what extent this differs from other sites in the organism where striated muscles play this role (Gopalakrishnan et al., 2015; Zhao and Dhoot, 2000).

Perturbations of esophagus function leads to dysphagia and other physiopathological disorders that impair swallowing and transfer of bolus to the stomach (Sheehan, 2008). ESM share a common origin with branchiomeric head muscles (Gopalakrishnan et al., 2015; Heude et al., 2018) and like those muscles, *Tbx1* and *Isl1* act as upstream regulators of ESM development (Gopalakrishnan et al., 2015). In *Tbx1-null* embryos, Isl1-derived myogenic cells fail to seed the anterior esophagus, suggesting that *Tbx1* acts genetically upstream of *Isl1* in ESM progenitors (Gopalakrishnan et al., 2015). Initially, CPM-derived progenitors are seeded at the bottom of the oropharyngeal cavity by E13.5. Then, Isl1-derived ESM progenitors colonize the esophagus by migration and differentiation until the third week of perinatal growth (Gopalakrishnan et al., 2015; Romer et al., 2013). How these *Isl1* progenitors colonize the structure while restricting premature differentiation remains unknown.

Muscle progenitors undergo short-range displacement or long-range migration for establishing skeletal muscles, as exemplified by myotomes and limbs, respectively. Progenitors originating from ventral somites delaminate and emigrate to distal sites to give rise to trunk, limb and tongue muscles (Bladt et al., 1995; Brand-Saberi et al., 1996; Dietrich et al., 1999). This process is regulated by the tyrosine kinase receptor Met, expressed in migratory progenitors, and its ligand Scatter Factor/Hepatocyte Growth Factor (SF/HGF) expressed in mesenchymal cells along the migratory route (Bladt et al., 1995; Brand-Saberi et al., 1996; Dietrich et al., 1999). Knockout of either *Met* or *Hgf* in mice results in the absence of hypaxial muscles including limb muscles, diaphragm and the tip of the tongue (Bladt et al., 1995; Dietrich et al., 1999; Maina et al., 1996; Prunotto et al., 2004). Although second (hyoid) arch-derived muscles are affected in *Met* KO mice (Prunotto et al., 2004), a role for Met/HGF in establishing other CPM muscles including those in the larynx and esophagus has not been reported.

In the present study, we used mouse chimeras to circumvent lethality issues and assessed the cell-autonomous roles of *Tbx1* and *Isl1* in ESM progenitors. Using genetic loss-of-function and pharmacological inhibition approaches, we show that Met/HGF is critical for ESM patterning, but not for the establishment of adjacent laryngeal muscles. These studies unveil an unexpected *Tbx1/Isl1/Met* genetic hierarchy operating within a CPM-muscle group, thereby identifying distinct genetic signatures for these evolutionarily conserved mesodermal derivatives.

## RESULTS

### Requirement of *Tbx1* and *Isl1* in ESM specification

We showed previously that Tbx1-null embryos lack ESM, wherein Isl1-derived myogenic progenitors fail to colonise and pattern the esophagus (Gopalakrishnan et al., 2015). The absence of seeding of Isl1-derived ESM progenitors in the anterior esophagus of Tbx1-null mice could originate from cell-autonomous or non-autonomous defects. To distinguish between these possibilities, we generated two types of chimeric embryos to explore the epistatic relationship between *Tbx1* and *Isl1* during ESM formation. Embryonic chimeras are well-established tools that have provided key insights into the tissue-specific requirement of genes during mammalian development (Tam and Rossant, 2003).

We first generated chimeras by injection of *Isl1^lacZ^* (KI) ES cells (Sun et al., 2007) in *Tbx1*^-/-^ and control (*Tbx1*^+/-^) blastocysts to determine if *Tbx1/Isl1*-positive cells can colonize the esophagus in a *Tbx1* null environment. Here, β-galactosidase (β-gal) expression is under the control of the *Isl1* promoter to trace the ES-derived cells *in vivo* (Figure S1A,B). All Tbx1-null chimeric embryos analyzed between E14.5 and E15.5 lacked thymus glands (9/9) and 77% of them (7/9) were edemic and lacked the outer ear pinna indicating that the injection of *Isl1^lacZ^* ES cells (*Tbx1^wildtype^*) was not extensive enough to fully rescue the *Tbx1* knockout phenotype (Figure S1C,D). Analysis of embryos by whole mount X-gal staining showed that 5/5 chimeric *Tbx1*^-/-^ embryos contained β-gal^+^ cells in the esophagus (Figure S1E), though to variable extent in individual embryos when compared to heterozygous controls. Apart from its expression in ESM progenitors, *Isl1* is expressed in peripheral neurons (Pfaff et al., 1996) and in the pharyngeal and esophageal epithelium (Cai et al., 2003; Harel et al., 2009; Nathan et al., 2008). Therefore, we performed an analysis on tissue sections to assess β-gal expression at the cellular level. We observed that β-gal+ cells were present within the smooth muscle layers of the esophagus of Tbx1-null embryos (8/8 chimeras), and colocalised within Tnnt3+/Tuj1-(myogenic/non-neurogenic) cells (Figure S1F,G). While ESM colonization in chimeric *Tbx1*^-/-^ embryos appeared to be less efficient than in controls (determined by number of β-gal+ cells/section and Tnnt3+ muscle area/section), the relative number of β-gal+ cells/Tnnt3+ muscle area was non significantly altered (Figure S1H-J). Taken together, these data indicate that *Isl1^lacZ^* ES cells can colonize an overall *Tbx1*-null esophageal environment suggesting cell autonomous potential of Tbx1+/Isl1+ progenitors to seed and pattern the ESM. Of note, cells expressing lower levels of β-gal were present in the esophagus epithelia and connective tissue layers of both control and chimeric Tbx1-null embryos suggesting that in these cells expression from the endogenous *Isl1* locus was downregulated, and therefore the extent of the contribution of *Isl1^lacZ^* ES cells (*Tbx1^wildtype^*) cannot be unambiguously assessed. To circumvent this issue, we generated a second series of chimeras to address the intrinsic role of *Isl1* during foetal esophagus myogenesis.

To bypass the early embryonic lethality of Isl1-null embryos (Cai et al., 2003), we generated chimeric foetuses by injection of Isl1-null ES cells into wildtype (WT) mouse blastocysts. We targeted Isl1-null and control (WT) ES cells with a constitutively expressing *lacZ* cassette (*pCAG-nlacZ*; nuclear β-gal activity) to trace ES cell derivatives ubiquitously and independently of *Isl1* expression (Figure 1A-B). Macroscopic examination of chimeras at E16.5 did not reveal obvious developmental defects in Isl1-null chimeras compared to controls. Immunostainings on sections were then performed to evaluate the contribution of Isl1-null (ES:*Isl1*^-/-^; *nlacZ*) and control (ES:WT; *nlacZ*) β-gal^+^ ES-derived cells to the esophagus myogenic population. For reference, contribution of β-gal^+^ ES-derived cells was compared with Isl1 lineage tracing (*Isl1*^*Cre*/+^*;R26*^*mT/mG/*+^ embryos), whereby GFP+ Isl1-derived CPM cells contribute to the esophagus myogenic population (Myod/Myog/Tnnt3+), but not to the esophagus smooth muscle (SMA+) and striated muscle of tongue that develop in an *Isl1*-independent context (Figure 1C-E). We then quantified the amount of chimerism and β-gal+ cells in the esophagus SMA+ and Myod/Myog+ populations and in myogenic cells of the tongue (n=3, Figure 1F-K). In both Isl1-null and control chimeras, the overall percentage of chimerism in the tongue was similar to that observed in the muscularized layers of the esophagus (Figure 1L-M, left pannels). In the esophagus of control chimeras, β-gal+ gave rise to both SMA+ (31%-46%) and Myod/Myog+ populations (25%-43%) (Figure 1F,G,L). In contrast, *Isl1*-null/β-gal+ cells were excluded from the esophagus Myod/Myog+ cells (Figure 1I,J,L), whereas they contributed to a similar extent to esophagus smooth muscle and tongue myogenic cells in both *Isl1*-null and control chimeras. These results show that *Isl1* is necessary cell-autonomously for progenitor cells to adopt a myogenic cell fate in the esophagus (Figure 1B).

**Figure 1.**
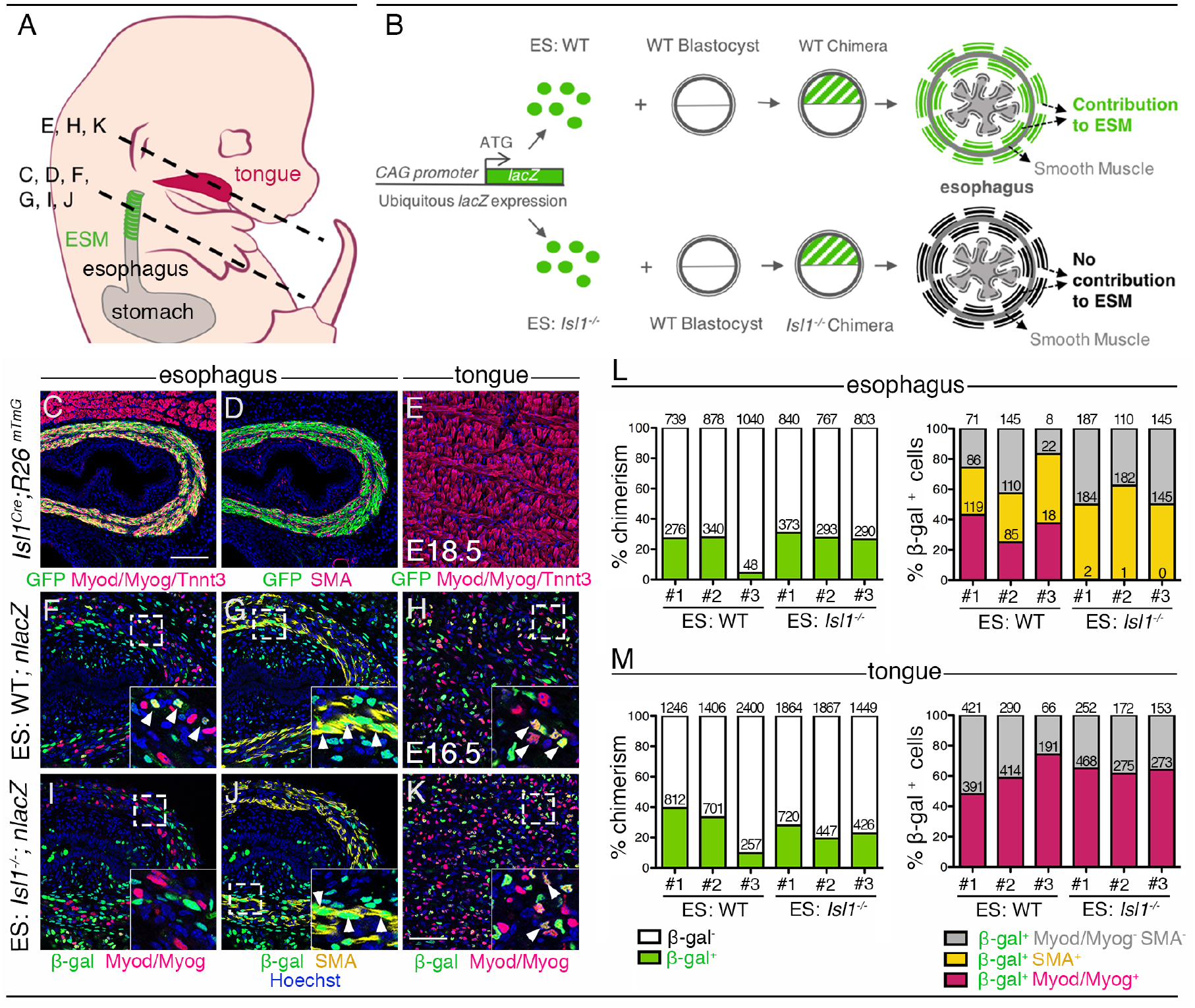
Cell-autonomous role of *Isl1* in esophagus myogenic progenitors. (A) Structures and levels analysed in the study. (B) Schematic summary of the chimera experiment. (C-E) Immunostainings on coronal cryosections of a E18.5 *Isl1^Cre^;R26^mTmG^* mouse for the GFP reporter, Myod/Myog/Tnnt3 (myogenic markers) and SMA (smooth muscle actin) in the esophagus and tongue. Note that *Isl1*-derivatives include the esophagus striated muscle but not the esophagus smooth muscle layers and tongue muscle (n=2). (F-K) Immunostainings on coronal cryosections of E16.5 WT (ES:WT;*nlacZ*) and *Isl1*^-/-^ (ES: *Isl1^-/-^;nlacZ*) chimeras for the β-gal reporter, Myod/Myog (myogenic markers) and SMA (smooth muscle actin) in the esophagus and tongue (n=3 each condition). Insets (bottom, right), higher magnifications. White arrowheads indicate examples of β-gal colocalisation with the myogenic markers. (L-M) Percentage of chimerism and of β-gal+ cell contribution to myogenic populations in the esophagus and tongue of WT (ES:WT) and *Isl1*^-/-^ (ES: *Isl1*^-/-^) chimeras (n=3 each condition, #1-3; three different section levels scored). The number of cells counted on 3 different section levels are reported in columns. Note that the *Isl1*^-/-^ ES-derived cells do not form ESM progenitors but contribute to both esophagus smooth muscle layers and tongue. Scale bars: A, 100 μm; I, 50 μm.

### Spatiotemporal activation of the ESM myogenic program

ESM development is biphasic, with initial seeding of *Isl1*-positive myogenic progenitors at the anterior esophagus followed by anterior-posterior migration and differentiation. To determine the expression of *Isl1* relative to the commitment MRF genes during ESM patterning, we performed RT-qPCR analysis at key stages of ESM development: at E15.5 and E17.5 when one third and two thirds of the esophagus contain ESM progenitors, respectively; then at 3 weeks postnatally when the entire esophagus is muscularized. *Isl1* and *Myf5* were detected in the anterior, middle and posterior esophagus as ESM progenitors colonize the structure from E15.5 to 3 weeks postnatally (Figure 2A-C). *Isl1* expression was also detected in the stomach, as already described for the gastric epithelium (Das and May, 2011). We next performed RT-qPCR analysis for *Isl1* in myogenic cells isolated from *Tg:Pax7nGFP* mice where Pax7+ progenitors can be isolated from mid-embryonic stages (Sambasivan et al., 2009) (Figure 2D). *Isl1* was expressed in Pax7-nGFP+ esophagus progenitors, whereas expression was low or undetectable in those isolated from the masseter at all stages analyzed. Therefore, *Isl1* expression is maintained in myogenic progenitors throughout ESM development.

**Figure 2.**
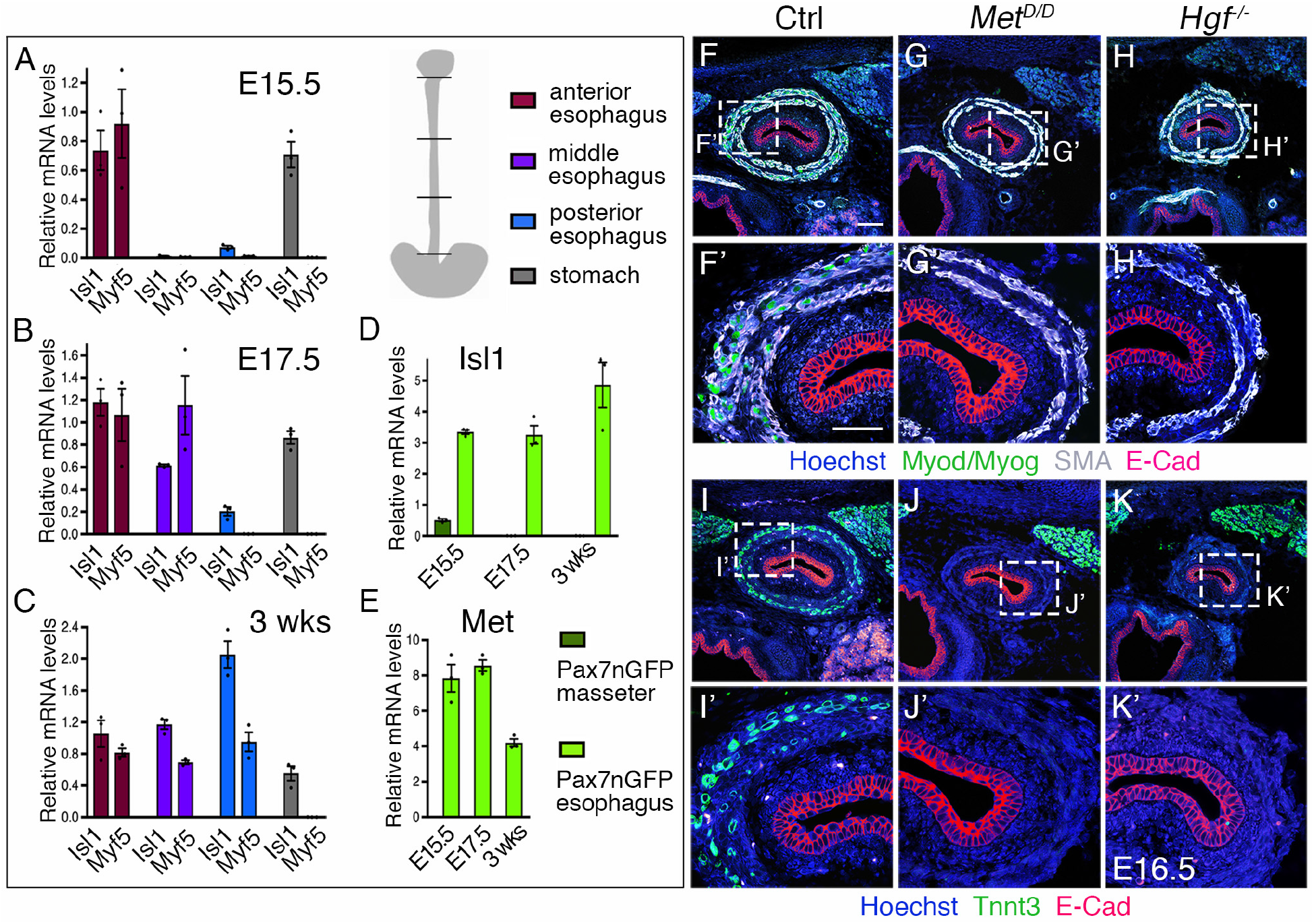
Regulation of esophagus striated muscle patterning involves Met/HGF signalling. (A-C) RT-qPCR analysis for *Isl1* and *Myf5* at E15.5 (A), E17.5 (B) and 3 weeks postnatal (C) in different esophagus portions and stomach indicated in the schematic view (top, right). The low level of *Isl1* expression in the posterior esophagus at foetal stages might reflect contamination from the anterior stomach at the esophagus interface. (D-E) RT-qPCR analysis for *Isl1* and *Met* at E15.5, E17.5 and 3 weeks postnatal in *Tg:Pax7nGFP* cells isolated by FACS from the masseter or esophagus. All data points are plotted and presented as the mean ± SEM (error bars) (n=3 each condition) (F-H) Immunostainings on coronal cryosections of E16.5 control, *Met^D/D^* and *Hgf^D/D^* foetuses for Myod/Myog (myogenic progenitors) and SMA (smooth muscle actin). E-Cad labels the esophagus lumen epithelium. Higher magnifications are shown in (F’-H’). (I-K) Immunostainings on coronal cryosections of E16.5 control, *Met^D/D^* and *Hgf*^-/-^ foetuses for the myofibre marker Tnnt3 and E-Cad. Higher magnifications are shown in (I’-K’). Note the absence of ESM formation in both *Met* and *Hgf* mutants (n=3 each condition). Scale bars: F, 100 μm; F’, 50 μm.

We then asked what molecular pathways would guide ESM progenitors to undergo A-P migration. Given the key role for Met/HGF signalling during myogenic migration and cell proliferation (Bladt et al., 1995; Dietrich et al., 1999; Maina et al., 1996; Prunotto et al., 2004), we performed RT-qPCR analysis for *Met* in Pax7nGFP+ in the esophagus. Notably, ESM progenitors showed transcript abundance of *Met* at fetal stages and lower expression levels postnatally when ESM colonization was complete (Figure 2E) suggesting that Met/HGF signalling might be implicated in A-P migration of ESM progenitors.

### Severe loss of ESM in *Met and Hgf* mutants

To address the role of Met/HGF signalling during ESM formation, we examined *Met^D/D^* and *Hgf*-null mutants (Maina et al., 1996; Schmidt et al., 1995). We first analysed the esophagus phenotype of *Met* and *Hgf* mutants at E16.5 by immunostainings on tissue sections for early myogenic and myofiber markers (Myod/Myog/Tnnt3), for smooth muscle (SMA) and lumen epithelium (E-Cad) markers. Interestingly, *Met^D/D^* and *Hgf*-null foetuses showed absence of striated muscles in the esophagus, while the smooth muscle layers and lumen epithelium appeared unaffected (Figure 2F-K). As expected, these mutants lacked limb muscles typical of the *Met^D/D^* and *Hgf*-null phenotypes (Figure S2A-F). However, RT-qPCR analysis in the esophagus and limb of *Met* mutants at E15.5 revealed a decrease but not loss of *Isl1* and *Myf5* expression compared to absence of *Pax7* and *Myf5* observed at limb level (Figure S2G).

Given this observation, we investigated whether myogenic cells are present at the anterior-most part of the esophagus and adjacent *Isl1*-derived muscles in the *Met^D/D^* foetuses (Figure 3A-C). Analysis on sections revealed that Isl1+ progenitors had seeded the anterior esophagus smooth muscle layers in mutant embryos similarly to controls at E13.5 (Figure 3A,B,B’). Moreover, analysis of *Met*^*D*/+^; *Myf5*^*nlacZ*/+^ (control) and *Met^D/D^; Myf5*^*nlacZ*/+^ (mutant) esophagi at E15.5 (Figure S3A,B) and E17.5 (Figure 3D,E) showed that Myf5+ myogenic cells were also present in the anterior-most portion of the esophagus in the mutant, whereas in the controls, colonization had proceeded posteriorly. Of note, the neuronal and smooth muscle lineages appeared to be present and patterned in the mutant (Figure 3D,E).

**Figure 3.**
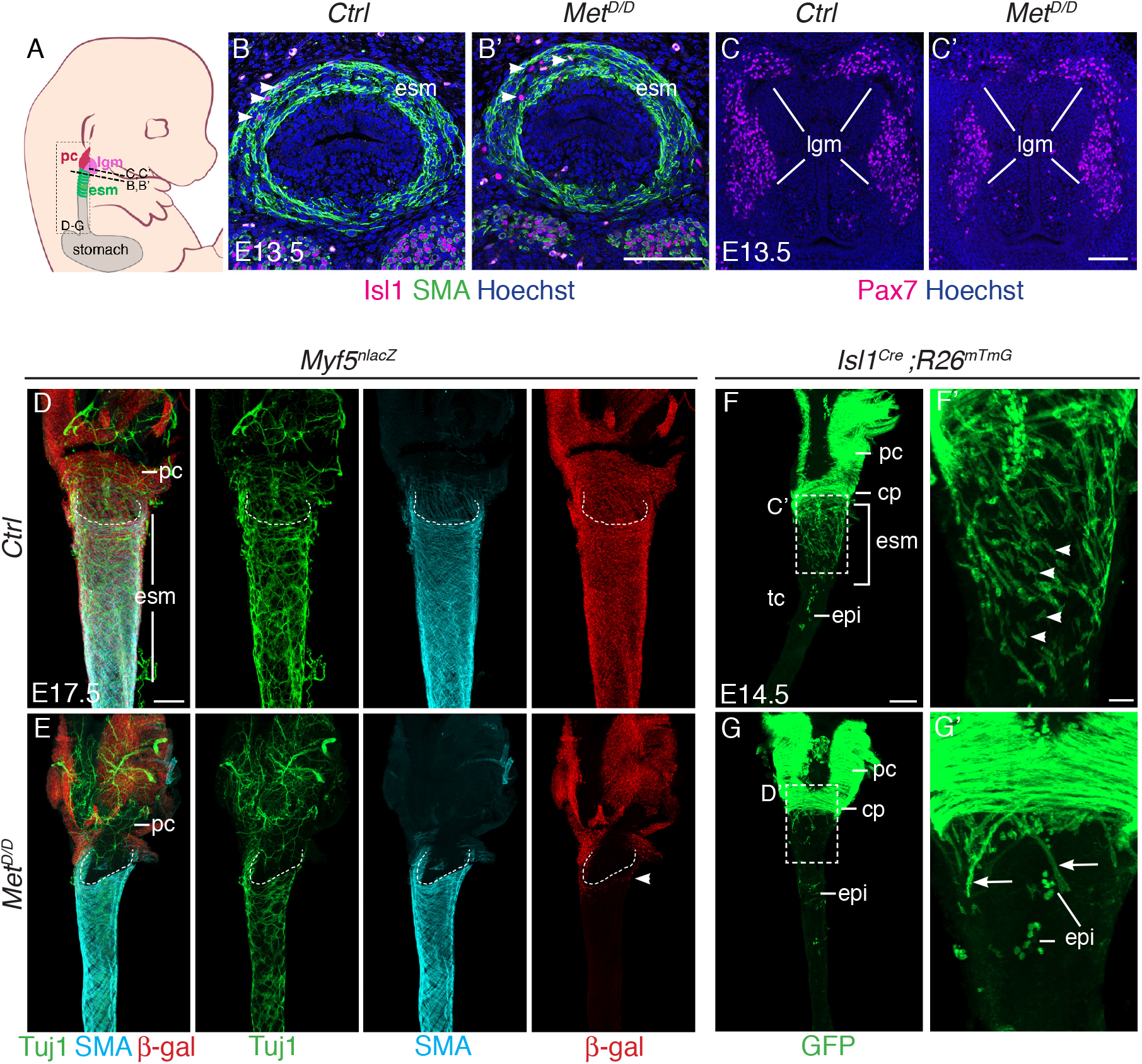
Isl1 progenitors are present anteriorly in the esophagus of Met^D/D^ mutants. (A) Structures and levels analysed in the study. (B,B’) Immunostainings on coronal cryosections of E13.5 control and *Met^D/D^* embryos for *Isl1* expressing progenitors (white arrowheads) and smooth muscle cells (SMA) in the esophagus (n=2). (C,C’) Immunostainings on coronal cryosections at the laryngeal level of E13.5 control and *Met^D/D^* embryos for *Pax7* (D-G) Whole mount immunostaining of the upper esophagus of Met mutant and control embryos. (D,E) Ventral views of E17.5 esophagi stained for Tuj1 (neurons), SMA (smooth muscle actin) and β-gal (Myf5 expressing progenitors). White dotted lines outline the shape of the esophagus entry. White arrowheads point to *Myf5*^*nlacZ*+^ progenitor cells present in the upper esophagus in the mutant. (F,G) Dorsal views of E14.5 stained for GFP (*Isl1* lineage tracing). *Isl1*-derived muscle progenitors remain largely as mononucleated cells (F,F’, white arrowheads) while GFP+ fibers are mostly seen in the anterior esophagus of the mutant (G,G’, arrows). cp, cricopharyngeous muscle; epi, epithelial *Isl1*-derived cells; esm, esophagus striated muscle; lc, laryngeal cecum; lgc, laryngeal cartilages, lgm, laryngeal muscles; pc, pharyngeal constrictor. Scale bars: B’, C’, F’, 50 μm; D, F, 200 μm

Next, we investigated the fate of the sporadic myogenic cells remaining in the anterior esophagus of *Met*-null foetuses. To this end, we combined the *Met^D/D^* mutant with *Isl1* lineage tracing. Analysis of E14.5 and E17.5 *Met*^*D*/+^; *Isl1*^*Cre*/+^ *;R26*^*mT/mG*/+^ control embryos showed that mGFP+ mononucleated cells could be abundantly detected between myofibres (Figure 3F, Figure S3C). However, very few mononucleated mGFP+ cells were detected between the residual myofibers in the anterior-most part of the esophagus in the *Met* mutant (Figure 3G; Figure S3D). Strikingly, the adjacent *Isl1*-derived laryngeal and pharyngeal muscles were unaffected in the Met-null foetuses (Figure 3C,C’,D-G; Figure S3A-D). Therefore, these observations indicate that Met/HGF affect only a subset of posterior CPM-derived progenitors that are critical for colonisation of the esophagus but not for the development of adjacent *Isl1*-lineage derived muscles.

Taken together, these results indicate that *Met* acts downstream of *Isl1* in the molecular hierarchy of ESM formation and that Met/HGF signalling is not implicated in initial seeding of *Isl1*-derived progenitors at the anterior esophagus, but rather during the second phase of migration within this structure.

### Requirement of Met/HGF signalling for A-P migration of ESM progenitors

We then wondered if the defect in ESM formation in *Met* mutants is due to an increase cell death or deficient migration of *Isl1*^+^ progenitors. We first tested if *Isl1*-derived ESM progenitors undergo apoptosis in the *Met^D^* early embryos. TUNEL analysis of E13.5 mutants showed that Isl1+ progenitors are not apoptotic (Figure S4A,A’).

Next, we investigated the role of Met in the migration of mGFP+ cells in an *ex vivo* esophagus explant culture system once they had colonized the upper esophagus (Figure 4A). To this end, we employed static and time-lapse confocal microscopy in combination with two selective ATP-competitive inhibitors of Met, PF-0417903 and MGC-265 or DMSO as control, on E14.5 *Isl1^Cre^;R26^mT/mG^* esophagus (Figure 4B-D). On static cultures (followed up to 24h) and time lapse imaging (up to 14h), we observed a mGFP+ mononucleated cell front that remained throughout the entire length in control cultures (Figure S4B, Figure 4B, Movie S1). In addition, time-lapse movies showed that mGFP+ cells explored the esophagus scaffold repeatedly changing their direction of migration, but had a net movement posteriorly towards the stomach (Figure 4E, Movie S1). Upstream of the mononucleated cell front, mGFP+ cells also migrated posteriorly in between forming fibers (Movie S1). In contrast, upon addition of Met inhibitors, mGFP+ cells progressed less towards the posterior end (Figure 4C,D, Movie S2,S3) and had shorter cell trajectories (Figure 4F-H). Quanfication of migration parameters revealed that in presence of Met inhibitors, mGFP+ cells had a reduced velocity, efficiency, and net displacement towards the posterior end when compared to control cells (Figure 4I-L). Interestingly, mGFP+ fibers appeared rapidly in the inhibitor treated cultures in positions where the cell density appeared higher (Figure 4C, D, Figure S4C), a phenotype that resembled *Met*-null embryos (Figure 4G, Figure S3D). To confirm this, we examined the proliferation and differentiation status of ESM myogenic cells *in vivo* by EdU labelling. Analysis of E14.5 and E15.5 embryos showed that myogenic cells in *Met^D^* mutant embryos have a proliferation rate that is one third of controls (45,6% for Ctrl; 12,5% for Mutant at E14.5, Figure S4D) and a higher predisposition to differentiation as assessed by Myogenin expression (33% for Ctrl; 66% for Mutant at E14.5, Figure S4E). In summary, our *ex vivo* and *in vivo* analysis support the notion that Met/HGF signalling is required for A-P migration of ESM progenitors once they have colonized the upper esophagus, and this effect is accompanied by an apparent precocious differentiation of *Isl1*-derived progenitors.

**Figure 4.**
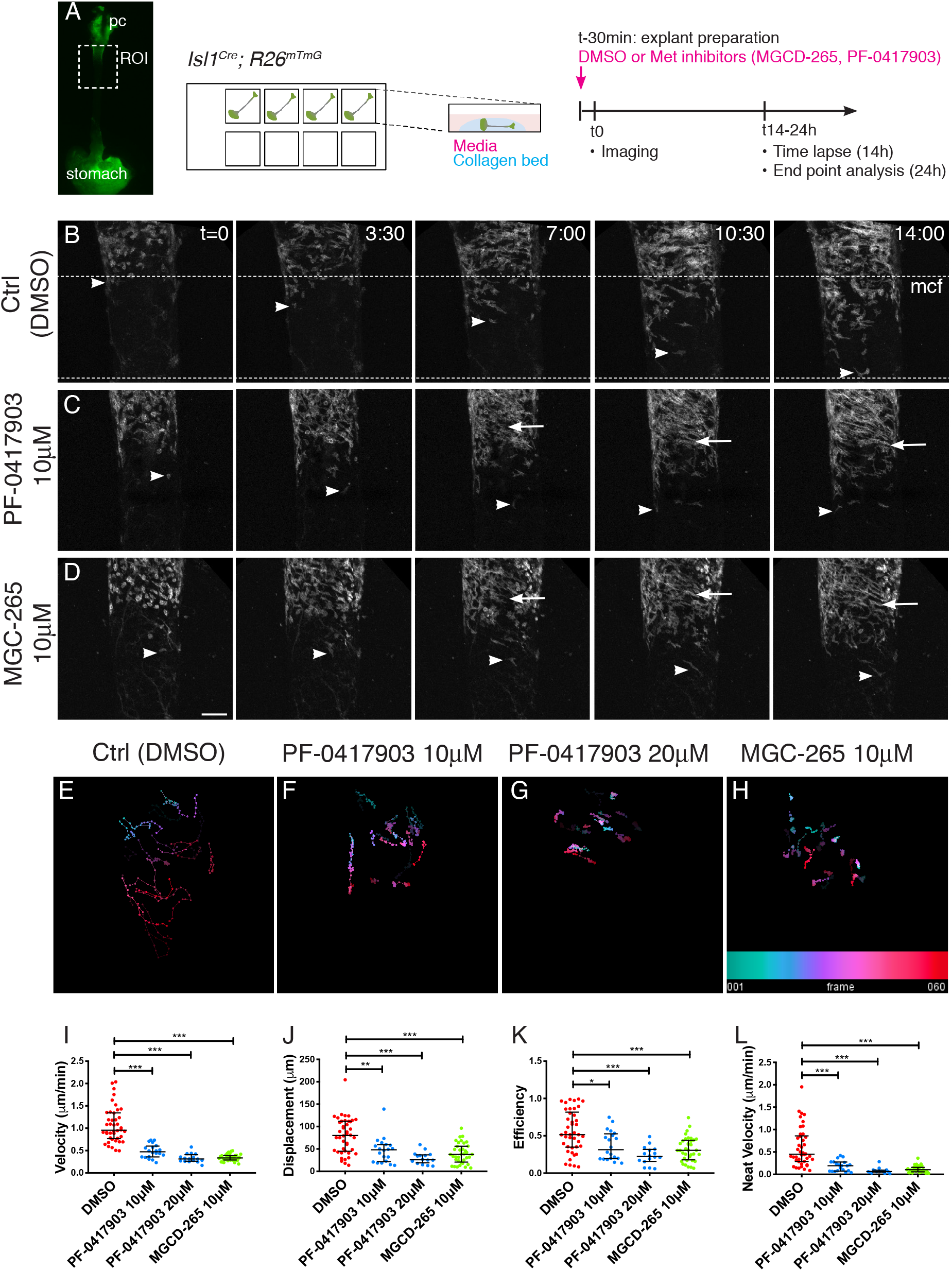
Met/HGF signalling is required for migration of *Isl1*-derived myogenic progenitors. (A) Macroscopic view of *Isl1^Cre^;R26^mTmG^* E14.5 dissected esopaghus used for explant culture and live imaging. Esophagi are placed in collagen beds in individual Ibidi wells. Met inhibitors (MGCD-265, PF-0417903) or control (ctrl, DMSO) is added to explants 30min before imaging. Explants are kept for 14h for live imaging (with an image taken every 12-15min) or 24h for analysis at fixed time points (Suppl. Figure 4). (B-D) Maximum projection of time series from a time-lapse experiment of esophagi explant culture in the presence of DMSO (B), 10μM PF-0417903 (C) or 10μM MGCD-265 (D). White arrowheads point to *Isl1*-derived progenitor cells present at the mononucleated cell front (mcf). White arrows highlight the high numbers of fibers that appear progresively in the inhibitor condition. Time (t) is indicated in hours. Dotted lines show the overall advancement of the mcf in the control condition. (E-H) Temporal color coded 2D images of GFP+ cell trajectories tracked in the time lapse movies in control and inhibitor treated explant cultures (related to Movies S1, S2, S3). (I-L) Quantification of cell velocity (in μm/min; I), displacement (μm, the lenght of the resultant vector between ti and tf of the track, J), efficiency (ratio between the displacement and the distance covered by the whole track, K), net velocity (μm/min, ratio between the displacement and total time of the track) in control and inhibitor treated explant cultures. Dots, individual cells tracked (from n=2 experiments). Mean ± SEM. Statistical significance was assessed by a Mann-Whitney test. pc, pharyngeal constrictor. Scale bar: D, 100μm

### Single cell analysis of ESM progenitors defines relationships between *Met* and *Isl1*

We then decided to examine in detail the expression pattern of *Hgf* and *Met* in relation to *Isl1*-derived progenitors. Owing to the limited diffusion efficiency of HGF *in vivo*, both receptor and ligand expressing cells are expected to be found in close proximity to each other (Dietrich et al., 1999). In situ hybridization (Rnascope®) on E14.5 *Isl1^Cre^;R26^mT/mG^* embryo cryosections, revealed that *Hgf* is expressed adjacent to mGFP+ cells in a bilayered concentric pattern corresponding to the smooth muscle layers of the esophagus (Figure 5A,B). As expected, Met was expressed at high levels in *Isl1*-derived mGFP+ cells, and also in luminal epithelial cells (Figure 5C). However, the levels of Met transcript in mGFP+ cells were heterogeneous. Co-immunostaining with Myod and Myog antibodies to detect differentiating myogenic cells, revealed that 44% of mGFP+ cells were Myod+/Myog+ and that 80% of these Myod+/Myog+ cells had low levels of *Met* transcript (score 0/1; Figure 5C1,C2,D,E). Conversely, 90% of Myod-/Myog-cells, expressed high levels of *Met* (score 3/4) (Figure 5C1,C2,E). Therefore, expression of *Met* was inversely correlated with the differentiation status of *Isl1*-derived cells.

**Figure 5.**
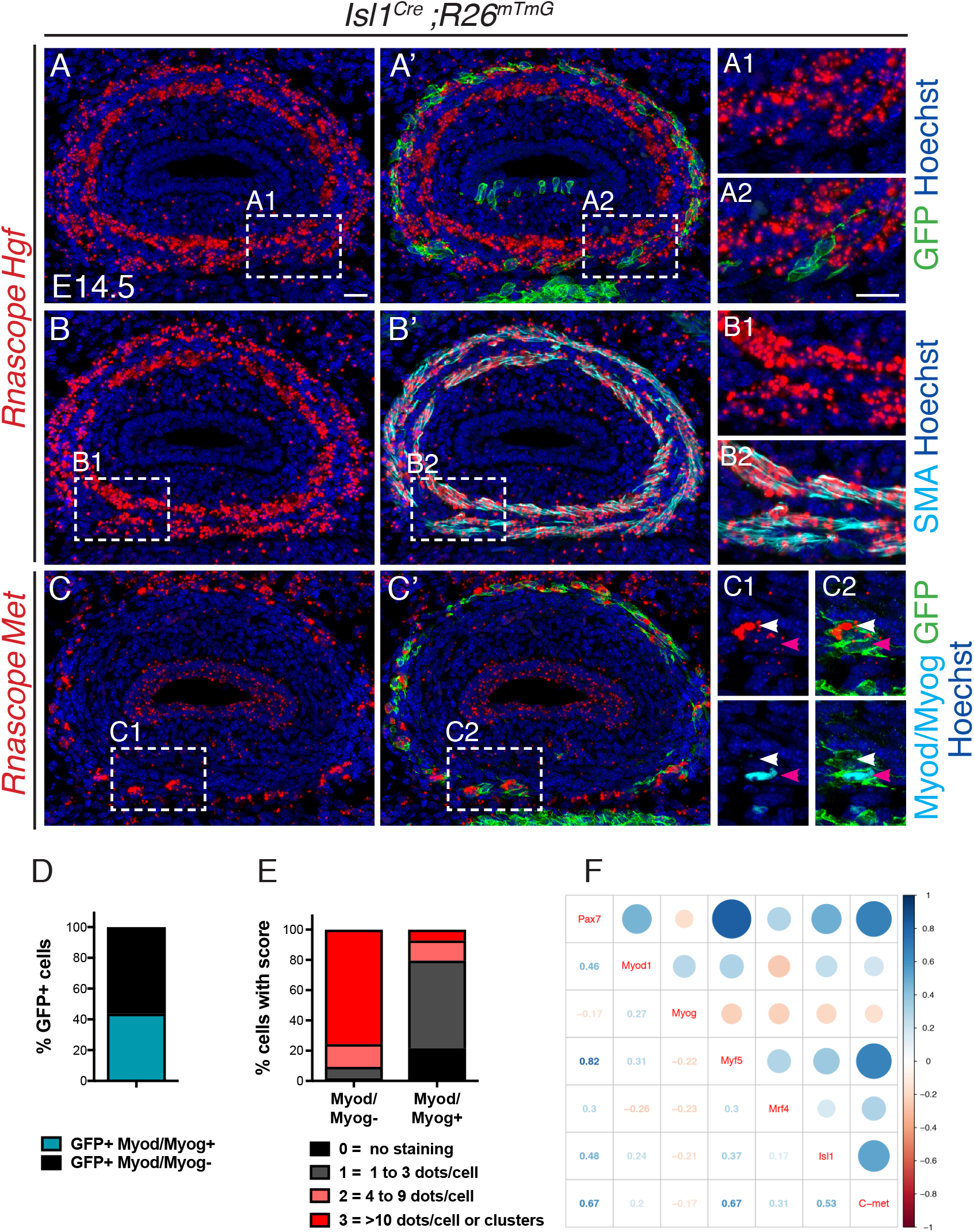
*Met* is expressed in undifferentiated *Isl1*-derived myogenic progenitors. (A-C) In situ hybridization on transverse cryosections at the esophagus level of E14.5 *Isl1^Cre^;R26^mTmG^* embryos for *Hgf* (A, B) and *Met* (C), combined with immunofluorescence for GFP (Isl1-derived progenitors), SMA (smooth muscle actin) and Myod/Myog (myogenic cells). Note that *Hgf* is expressed adjacent to *Isl1*-derived cells (A1,A2) by SMA+ cells (B1,B2). Met is expressed by *Isl1*-derived cells (C1,C2, top panels) but at levels inversely correlated to Myod/Myog+ expression (C1, C2, bottom pannels). (D) Histograms of the percentage of Myod/Myod- and Myod/Myog+ *Isl1*-derived GFP+ cells. (E) Histograms of the percentage of cells in (D) with a defined Rnascope® score for Met expression. (F) Correlogram. The upper part of the mixed correlogram displays graphically the degree of relationship between genes with the bigger the circle, the higher the Spearman’s correlation coefficient, the redder, the more negative, the bluer, the more positive. Significant correlations are highlighted with an orange background. The lower part shows the values of the Spearman’s coefficient, and the p-values (between brackets) for the significant ones. Scale bars: A, A2, 20μm

To investigate the relative expression status of these myogenic markers in more detail, we performed single cell RT-qPCR analysis of *Isl1*-derived ESM progenitors. The mononucleated cell front in the esophagus of E15.5 *Isl1^Cre^;R26^mT/mG^* mice was dissected and the expression of myogenic markers was examined in GFP+ isolated by fluorescence activated cell sorting (FACS) (Figure S5A). The normalized relative expression of the studied genes for all the filtered cells that was annotated in a heatmap (Figure S5B) revealed two groups of genes. The first group included *Isl1, Met, Pax7* and *Myf5* which were detected in nearly all the single cells analysed. The second group included *Mrf4, Myog* and *Myod1* which were expressed in a subset of cells. To assess the degree of relatedness between genes, we calculated the Spearman’s correlation between all pairs of genes and noted that the expression of *Isl1, Met, Pax7* and *Myf5* was significantly positively correlated (Figure 5F). In contrast, *Isl1* expression showed no significant correlation with the expression of more downstream MRF genes (Myod, Myog, and Mrf4). This indicates, that *Isl1* is associated with the upstream state as expected, and that *Isl1* and *Met* likely act concomittantly in myogenic progenitors during ESM formation.

## DISCUSSION

In vertebrates, branchiomeric head and neck muscles share a common CPM progenitor pool regulated by upstream molecular players including *Tbx1* and *Isl1*. Here, we uncover a cell-autonomous requirement of *Tbx1* and *Isl1* in the specification of CPM-derived esophagus myogenic progenitors. In addition, we show for the first time a unique dependency of myogenic progenitors on Met/HGF signalling pathway for esophagus spatio-temporal patterning. Surprisingly, laryngeal muscles that also originate from the posterior pharyngeal arches, are unaffected in *Met* mutants, thereby uncoupling the genetic requirements between ontogenically similar groups of CPM-derived muscles. These findings highlight distinct genetic hierarchies operating with CPM derivatives, and provide a framework to address myopathies of cranial origin (Figure 6).

**Figure 6.**
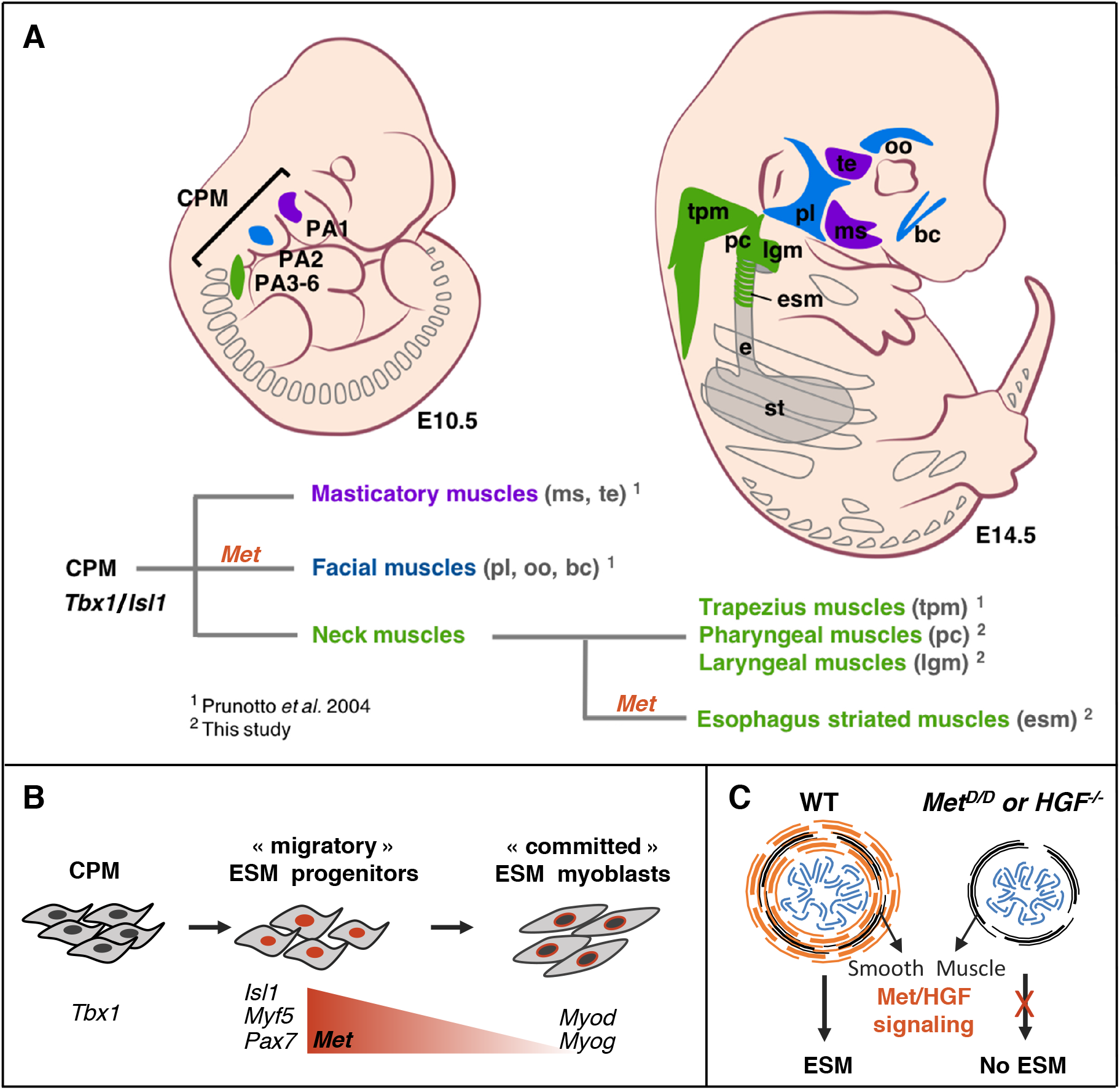
*Tbx1-Isl1-Met* genetic pathway regulates only a subset of CPM-derived muscles. A) Masticatory (purple) and facial (blue) muscles originating from anterior pharyngeal arches (PA1-2) are indicated. Neck muscles (green) derived from posterior PAs including trapezius, pharyngeal and laryngeal muscles, develop in a *Met*-independent context, while esophagus striated muscles are under the control of Met/HGF signalling. B) A *Tbx1/Islet1/Met* genetic hierarchy acts in uncommited ESM progenitors. Then, *Met* expression decreases in myoblasts during myogenic commitment. C) Absence of ESM formation in the *Met* and *HGF* mutants. CPM, cardiopharyngeal mesoderm; bc, buccinator; e, esophagus; esm, esophagus striated muscles; lgm, laryngeal muscles; ms, masseter; oo, orbicularis oculi; PA1-6, pharyngeal arches 1-6; pc, pharyngeal constrictor; pl, platysma; st, stomach; te, temporal; tpm, trapezius muscles.

### Cell-autonomous role of *Isl1* during esophagus myogenesis

Recent genetic studies revealed that neck muscles including pharyngeal and laryngeal muscles, the trapezius and the esophagus originate from posterior pharyngeal arch mesoderm derived from an *Isl1*-lineage (Gopalakrishnan et al., 2015; Heude et al., 2018; Lescroart et al., 2015; Tabler et al., 2017). We previously demonstrated that *Tbx1* and *Isl1* genes play key upstream roles during ESM formation (Gopalakrishnan et al., 2015). In Tbx1-null embryos, *Isl1*-derived ESM fail to form, indicating that *Tbx1* acts upstream of *Isl1* during esophagus myogenesis (Gopalakrishnan et al., 2015). Given that Isl1 promotes cell proliferation and represses myogenic differentiation, Isl1 has been proposed to exert a conserved role in the specification of CPM progenitors (Cai et al., 2003; Diogo et al., 2015; Harel et al., 2009). However, the intrinsic role of Isl1 in CPM derivatives has not been addressed due to early embryonic lethality (Cai et al., 2003). Here, by means of chimeric analysis, we show that *Isl1*-null ES cells are specifically excluded from the ESM indicating that *Isl1* acts cell-autonomously during ESM formation at fetal stages, further supporting its role in the specification of branchiomeric myogenic progenitors.

### Met/HGF signalling drives esophagus progenitor migration

During development, positional information that includes migration cues is often imparted to cells through intercellular signalling to allow proper spatio-temporal patterning. Several studies have uncovered the role Met receptor and its ligand HGF in the proliferation and motility of long-range myogenic progenitors (Trusolino et al., 2010). The *Met* allele used here (*Met^D^*) carries a mutation in two phosphotyrosines (Tyr1349, Tyr1356) in the carboxy-terminal tail, which completely abrogates Met function and recapitulates the *Met* null phenotype (Bladt et al., 1995; Maina et al., 1996; Maina et al., 2001). Previous work showed that in *Met* and *Hgf* mutants, Pax3-derived hypaxial muscles are missing, while other trunk muscle groups appear unaffected (Bladt et al., 1995; Dietrich et al., 1999; Maina et al., 1996; Prunotto et al., 2004).

Here, we show that ESM is also absent in both mutants. At early stages, *Isl1*+ progenitors seeded in the anteriormost part of the esophagus are not apoptotic in *Met^D/D^* mutants, suggesting that Met does not have a role in survival of premigratory myogenic cells. The first obvious deficiency observed in the *Met^D/D^* null is seen at the time Isl1+ progenitors colonize the smooth muscle scaffold (by E14.5). In the control, colonization progresses posteriorly, myofibers are formed while maintaining a pool of progenitor cells. In the mutant, only few *Isl1*-derived myofibers are present in the upper esophagus.,Therefore Met has a role in motility as well as the balance between proliferation and differentiation of *Isl1*-derived myogenic cells once they have entered the smooth muscle scaffold.

### A migration vs differentiation balance for esophagus myogenic progenitors

In the trunk, *Hgf* is first expressed adjacent to somites, and subsequently along the migratory route and at target sites in limb connective tissue (Dietrich et al., 1999). In the esophagus, we identified the smooth muscle layer, which serves as a scaffold for myogenic progenitor migration, to be a major source of *Hgf*. This finding underscores the functional relevance of smooth and striated muscle progenitor interactions for proper ESM patterning that is unique compared to muscle patterning elsewhere. *Hgf* expression at the time of myogenic progenitor migration was observed throughout the length of the esophagus, suggesting that Met/HGF signalling is important all along the migration phase. It has been proposed that a prolonged interaction between *Met* and *Hgf* may be required to prevent cell re-aggregation, thereby maintaining cell motility and prevent expression of the MRFs (Dietrich et al., 1999). Whether motility prevents or delays expression of the differentiation genes or differentiation gene expression stop motility is unclear. However, it has been shown that precocious expression of MRFs in dermomyotomal muscle progenitors prevents their migration into limb buds (Bonnet et al., 2010), while application of HGF results in reduction of *Myod* expression (Scaal et al., 1999).

Our pharmacological inhibition studies of Met receptor activity in esophagus explant cultures resulted in impaired progenitor cell migration and differentiation. Thus, Met/HGF signaling might have a role in maintenance of the undifferentiated state of migratory muscle progenitors. Accordingly, our in situ hybridization and single cell qPCR data showed that *Isl1* and *Met* act predominantly in uncommitted ESM progenitors (Pax7+, Myod/Myog-), and had decreased expression in committed cells. During limb muscle development, the Met receptor was reported to be under the direct transcriptional regulation of *Pax3* (Epstein et al., 1996). Interestingly, ESM development requires Met/HGF signalling in a Pax3-independent context. Hence the upstream modulator of c-Met expression in ESM progenitors remains an open question. *Pax3* and its paralogue *Pax7* have partially redundant functions in muscle progenitors (Relaix et al., 2006). Intringuingly, *Pax7* mutant mice have impaired ESM formation due to reduced proliferation and precocious differentiation at the migratory front (Chihara et al., 2015). Thus, it will be interesting to explore if Pax7 or Isl1 directly regulate *Met* expression in ESM.

### Myogenic diversity within CPM-derived muscles

An unexpected finding from our work is that CPM muscles originating from posterior pharyngeal arches are differentially affected in *Met* mutants. Head muscles derived from the second pharyngeal arch and giving rise to branchiomeric facial muscles (orbicularis occuli, buccinator, platysma) appear either strongly reduced or absent, while first arch derived masticatory (masseter, temporalis) and extraocular muscles are present in *Met^D/D^* mutants (Prunotto et al., 2004). However, we show that posterior branchiomeric neck muscles including pharyngeal and laryngeal muscles, are present in *Met^D/D^* mutants while adjacent ESM is absent. Thus, we have established a unique *Tbx1-Isl1-Met* genetic hierarchy in ESM progenitors that is distinct from other posterior branchiomeric muscles (Figure 6).

The genetic regulatory pathways that give rise to functionally distinct groups of muscles has provided critical information to understand heterogeneity in response to genetic diseases, such as DiGeorge syndrome where mutations in TBX1 result in the impared function of subsets of the craniofacial and pharyngeal apparatus with varied degrees of severity. Understanding the functional dynamics of *Tbx1* and *Isl1* in specific muscles groups will help uncover differences between the ontogenically similar group of CPM derived muscles. Uncoupling the genetic requirements of these distinct populations is necessary to provide a framework that will explain how human myopathies affect only subsets of muscles (Emery, 2002; Randolph and Pavlath, 2015).

## MATERIALS AND METHODS

### Animals

Animals were handled as per European Community guidelines and the ethics committee of the Institut Pasteur (CTEA) approved protocols. *Isl1^Cre^* (Srinivas et al., 2001), reporter mouse lines *R26R^mT/mG^* (Muzumdar et al., 2007), *Myf5^nLacZ^* (Tajbakhsh et al., 1996), *Tg: Pax7nGFP* (Sambasivan et al., 2009), and mutant mice carrying the *Tbx1^tm1pa^* allele (referred to as *Tbx1*^-/-^) (Jerome and Papaioannou, 2001), *Hgf* (Schmidt et al., 1995) and *Met* (referred as Met^D^) (Maina et al., 1996) mutant alleles were described previously. To generate experimental embryos for *Met^D/D^* together with *Isl1* and *Myf5* lineage tracings, *Met*^*D*/+^: *Isl1*^*Cre*/+^: *Myf5*^*nlacZ*/+^ males were crossed with *Met*^*D*/+^: *R26R^mTmG/mTmG^* females. Mice were kept on a mixed genetic background C57BL/6JRj and DBA/2JRj (Janvier Labs). Mouse embryos and foetuses were collected between embryonic day (E) E12.5 and E18.5, with noon on the day of the vaginal plug considered as E0.5.

### Generation of *Isl1*-null chimeras

For derivation of *Isl1-null* ES cells, males and females from *Isl1*^*Cre*/+^ genotype (Srinivas et al., 2001) were intercrossed to produce heterozygous and homozygous *Isl1-null* blastocysts. At E3.5, blastocysts were collected from uterine horns and put on culture for 3-6 days in ES derivation medium composed of GlutaMAX/DMEM (Gibco, 31966), 15% FBS (Biowest), 1% penicillin/streptomycin (Gibco, 15140, stock 100X), 1% sodium pyruvate (Gibco, 11360, stock 100mM), 0.1% β-mercaptoethanol (Gibco, 31350-010, stock 50mM), 1000U/ml ESGRO® recombinant mice leukemia inhibitory factor (LIF, Millipore, ESG1107, stock 10^7^U/ml) and 2i (1μM PD325; Axon Medchem 1408; and 1μM CH99; Axon Medchem 1386) on gelatin-coated wells with primary Mouse Embryonic Fibroblasts (MEFs). The disaggregation of ICM was performed with 5 min of 0.05% trypsin-EDTA (GIBCO, 25300-054) treatment and the cell suspension put on culture in ES derivation medium on MEFs. Derived ES cells were then expanded and genotyped by PCR with specific primers for amplification of *Isl1* WT and mutant sequences (WT primers: ccaagt gcagcataggcttcag; gcagaggccgcgctggatgcaagg, 230bp; Mutant primers: tcatgcaagctggtggctgg; gcagaggccgcgctggatgca agg, 633 bp).

To trace the ES clones, CAG-nlacZ and PGK-puro cassettes were cloned into a pBluescript to produce a *nlacZ* reporter puromycin resistant plasmid. Heterozygous and homozygous *Isl1-null;nLacZ* ES cells were electroporated (0.5-1 × 10^7^ cells) with 20 μg of linearized pCAGnlacZ-puro plasmid by using a BTX Harward apparatus ECM830 electroporator with one pulse at 240V for 15ms. Three days after transfection, positive clones were selected in ES derivation medium with puromycin (1.5μg/ml) for 5 days. ES colonies were picked into 24-well plates and tested for expression of the *nLacZ* reporter (X-Gal/immunostaining).

For chimera production, the β-gal+ selected clones were further expanded in ES culture medium (ES derivation medium without 2i) on MEFs. C57BL/6N females were superovulated and mated with C57BL/6N males. At E3.5, blastocyts were collected, injected with wildtype (control) or homozygous *Isl1*-null*;nlacZ* ES cells (2-6 cells/blastocyst) and were subsequently transfered into the uterus of 0.5 or 2.5 dpc pseudopregnant B6CBAF1 females (15-17 blastocyts/females). Chimeric fetuses were harvested at E16.5 or E18.5 for analysis. The collected fetuses were dissected in PBS at 4°C to remove the caudal part below the stomach, then fixed 3h at 4°C in 4% paraformaldehyde (PFA, Electron Microscopy Sciences, 15710), 0.5% Triton X-100 (SIGMA, T8787) and extensively washed in PBS with 0.1% Tween 20 (PBST) (SIGMA, P1379). To evaluate the contribution of ES cells to specimens, X-gal staining was performed on the dissected lower part of the fetuses. The analysis was performed by immunofluorescent stainings on cryosections of the rostral part of the fetuses.

### Generation of Tbx1-null chimeras

The Isl1 nuclear LacZ (nLacZ) knock-in mouse 129/SV ES line (*Isl1^lacZ^*) was obtained from Sylvia Evans (Sun et al., 2007). ES cells were cultured on MytomycinC treated embryonic primary fibroblasts onto gelatin coated dishes in DMEM-KO media (Gibco, 10829-018) containing 15% FBS (Biowest), 0,5% penicillin/streptomycin (Gibco, 15140, stock100x), 0.1% β-mercaptoethanol (SIGMA, M7522, stock 100mM in PBS), 1% L-Glutamine (Gibco, 25030024, stock 200mM) and 1000U/ml ESGRO® recombinant mice leukemia inhibitory factor (LIF, Millipore, ESG1107, stock 10^7^U/ml).

For ES cell injection and chimera production, 4 week old *Tbx1*^+/-^ females were superovulated and mated with *Tbx1*^+/-^ males (on a mixed genetic background C57BL/6JRj and DBA/2JRj, Janvier Labs). At E3.5, blastocyts were collected, injected with *Isl1^lacZ^* ES cells (6-12 cells/blastocyst) and were subsequently transfered into uteri of 0.5 or 2.5 dpc pseudopregnant B6CBAF1 females.

Chimeric fetuses were harvested at E14.5/E15.5 for analysis. The collected fetuses fixed 2h30 at 4°C in 4% paraformaldehyde 0.2% Triton X-100 and extensively washed in PBS at 4°C. For genotyping of the chimeric embryos, the visceral yolk sac layers were separated using the trypsin/pancreating method as described in (Manipulating the mouse embryo, Cold Spring Harbor Laboratory Press, 2003; Wallingford et al. Mech of Dev 2014). Briefly, yolk sacs were collected and incubated in Ca++/Mg++-free Tyrode Ringer’s saline solution containing 0.5% Trypsin (Gibco, 15090-046) and 2.5% Pancreatin (SIGMA, P-3292) for 4h at 4°C on individual wells of a 12 well plastic dish. Yolk sacs were then washed in GlutaMAX/DMEM (Gibco, 31966021) media buffered with 25mM HEPES (SIGMA, H0887) and then transferred into media contaning 10% FBS for at least 30 min at 4°C. The visceral endoderm (VEnd) and extraembryonic mesoderm (ExM) tissue layers of the visceral yolk sac were mechanically separated for genotyping. The VEnd layer is contributed exclusively by the host embryo while the ExM has dual contribution from ES cells and host embryo. DNA extraction was performed using ProteinaseK and PCR performed with the following primers: Tbx1_for: tgcatgccaaatgtttccctg, Tbx1_rs: gatagtctaggctccagtcca, Tbx1_rs_Neo: agggccagctcattcctcccac (WT band: 196bp; Mutant band: 450bp), lacZ_fw: atcctctgcatggtcaggtc, lacZ_rs: cgtggcctgattcattcccc.

For the analysis of lacZ+ chimeric embryos, the digestive tract including the pharynx, trachea, esophagus, heart, stomach and diaphragm was further dissected and X-Gal stained overnight at 37°C or embryos were processed for cryosections and immunostaining on sucrose/OCT as described above.

### Methods details

#### X-Gal staining and Immunofluorescense

Wholemount samples were analysed for β-galactosidase activity with 400 μg/ml X-Gal (SIGMA 15520-018; Stock solution 40mg/ml in DMSO) in PBS buffer containing 4 mM potassium ferricyanide, 4 mM potassium ferrocyanide, 0.02% NP-40 and 2 mM MgCl_2_ as previously described (Comai et al., 2014).

For immunostaining on cryosections, embryos and fetuses were fixed 3h in 4% PFA and 0,2-0,5% Triton X-100 at 4°C, washed overnight at 4°C in PBS, cryopreserved in 30% sucrose in PBS and embedded in OCT for cryosectioning. Cryosections (16-18μm) were allowed to dry for 30 min and washed in PBS. For immunostaining on paraffin sections, samples were fixed overnight in 4% PFA, dehydrated in graded ethanol series and penetrated with Histoclear II (HS-202, National Diagnostics) and embedded in paraffin. Paraffin blocks were sectioned at 12 μm using a Leica microtome. Sections were then deparaffinized and rehydrated by successive immersions in Histoclear, ethanol and PBS series. When needed, samples were then subjected to antigen retrieval with 10 mM Citrate buffer (pH 6.0) using a 2100 Retriever (Aptum Biologics).

Rehydrated sections were blocked for 1h in 10% normal goat serum, 3% BSA, 0.5% Triton X-100 in PBS. Primary antibodies were diluted in blocking solution and incubated overnight at 4°C. After 3 rounds of 15 min washes in PBST, secondary antibodies were incubated in blocking solution 1h at RT together with 1μg/ml Hoechst 33342 to visualize nuclei. Antibodies used in the study are listed in Table1. After 3 rounds of 15 min washes in PBST, slides were mounted in 70% glycerol in PBS for analysis. For Edu staining, immunostaining for primary and secondary antibodies was performed first, followed by the click chemical reaction using Alexa633 as a reactive fluorophore for EdU detection (Life Technologies C10350).

For whole mount immunostaining, embryos were fixed and washed as above. Esophagi were micro-dissected in PBST and incubated in blocking buffer (10% goat serum, 10% BSA, 0.5% TritonX-100 in 1X PBS) for 1h at RT in 2ml Eppendorff tubes. The tissue was then incubated with primary antibodies in the blocking buffer for 4-5 days at 4°C with rocking. The tissue was washed extensively for 2h-4h in PBST and then incubated in Fab’ secondary antibodies for 2 days at 4°C with rocking. The tissue was washed as above, dehydrated in 50% Methanol in PBS, 100% Methanol and then cleared with BABB and mounted for imaging as in (Yokomizo et al., 2012).

#### Rnascope® *in situ* hybridization

E14.5 embryos were collected, fixed overnight in 4% PFA, washed in PBS 3×15min, equilibrated in 15% and 30% sucrose and embedded in OCT. Tissue blocks were stored at −80C. 18μm thick cryosections were collected on Superfrost Plus slides and stored at −80 till use (less than 2 months).

RNAscope® probes Mm-Hgf (315631) and Mm-Met (405301) were designed commercially by the manufacturer and are available from Advanced Cell Diagnostics, Inc. In situ hybridization was performed using the RNAscope® 2.5 HD Reagent Kit-Red according to manufacturer’s instructions (Wang et al., 2012) with modifications. For sample pre-treatments: H2O2 treatment was 10min at RT, retrival was done for 2 min at 98°C and slides were digested with Protease Plus reagent for 15min at 40°C. The AMP1 to AMP6 steps were done as in the standard protocol. Before detection, samples were washed in PBS 3x 5min and immunostaining performed as above with fluorescent secondary antibodies. Sections were then washed in Rnascope® Wash buffer, detection done with Fast-Red A/B mix and slides mounted in Fluoromount-G (InterBioTech, FP-483331). As the Fast-Red chromogenic precipitate is also visible by fluorescence microscopy using the 555nm laser, sections were imaged using a 40x objective on a LSM700 microscope (Zeiss). For quantitation of Met RNAscope® staining, the number of individual signal dots or clusters per mGFP+ cell was counted manually on Fiji. Cells were attributed the score 1 (1 to 3 dots/cell), 2 (4 to 9 dots/cell) or 3 (more than 10 dots/cells or big clusters) and correlated to the presence or absence of Myod/Myog nuclear staining.

#### Enzymatic digestion for cell sorting

The masseter muscles and esophagi from *Tg:Pax7nGFP* timed embryos were dissected in cold PBS and kept in cold GlutaMAX/DMEM (Gibco, 31966) with 1% Penicillin–Streptomycin. For single cell qPCR analysis, only mGFP+ cells from the mononucleated cell front (mcf) of the esophagus of *Isl1^Cre^:R26^mTmG^* embryos were micro-dissected under a Zeiss SteREO Discovery V20 microscope. Samples were processed with enzymatic digestion mix containing 0.1% Trypsin (15090-046,Gibco®), 0,08% Collagenase D (Roche, 11088882001) and 10μg/ml of DNAse I (04536282001, Roche) in DMEM/Glutamax. Samples were incubated for 15min at 37°C under 300 rpm agitation, resuspended by gently pipetting up and down 10-15 times using a P1000 pipette, incubation and resuspension by pipetting were repeated for two additional 15 min enzymatic treatments. The digests were passed through a 70 micron then 40 micron SmartCell Strainers (Milteny Biotec) and digestion was stopped with Foetal Bovine Serum (FBS, Gibco). Cells were spun at 600g 15 min at 4°C and the pellets resuspended in 300μl of DMEM/2% FBS to be processed for FACS.

#### Quantitative RT-qPCR

Total RNA from esophagus portions and limbs was extracted through manual pestle tissue disruption in Trizol and purified with the Qiagen RNAeasy Mini purification Kit. Total RNA was extracted from *Tg:Pax7nGFP* cells isolated by FACS directly into cell lysis buffer (RLT) of the Qiagen RNAeasy Plus Micro purification Kit. cDNA was prepared from 0,4μg up to 5μg of total RNA by random-primed reverse transcription (SuperScript III, ThermoFisher 18010093) and real-time PCR was done using SYBR Green Universal Mix (Roche, 13608700) and StepOne-Plus Real Time PCR System (Applied Biosystems). TBP transcript levels were used for normalizations of each target (2ΔCT). At least three biological replicates and technical duplicates were used for each condition method (Schmittgen et al., 2008). For SYBR-Green, custom primers were designed using the Primer3Plus online software. Serial dilutions of total cDNA were used to calculate the amplification efficiency of each primer set according to the equation: E=10 – 1/slope. Primer sequences used are detailed in Table1.

#### Single-cell qPCR analysis

Gene expression in single cells was analysed using the Fluidigm Gene Expression Assay (BioMark). Briefly, oesophagus was dissected and digested with trypsin/collagenase to obtain a single cell suspension as described above. Single cells and bulk control (20 cells/well) were sorted directly on a FACS Aria III in 9 μl of Specific Target Amplification (STA) reaction mix from the CellsDirect One-Step qRT-PCR kit (Invitrogen) containing 0.2XTaqMan Gene Expression Assay mix. Pre-amplified cDNA (18 cycles) was obtained according to manufacturer’s note and was diluted 1:5 in TE buffer for qPCR. Multiplex qPCR was performed using the microfluidics Biomark system on a Biomark HD for 40 cycles. The same TaqMan probes were used for both RT/STA and qPCR. TaqMan assays used in the study are listed in Table 2.

**Table1.**
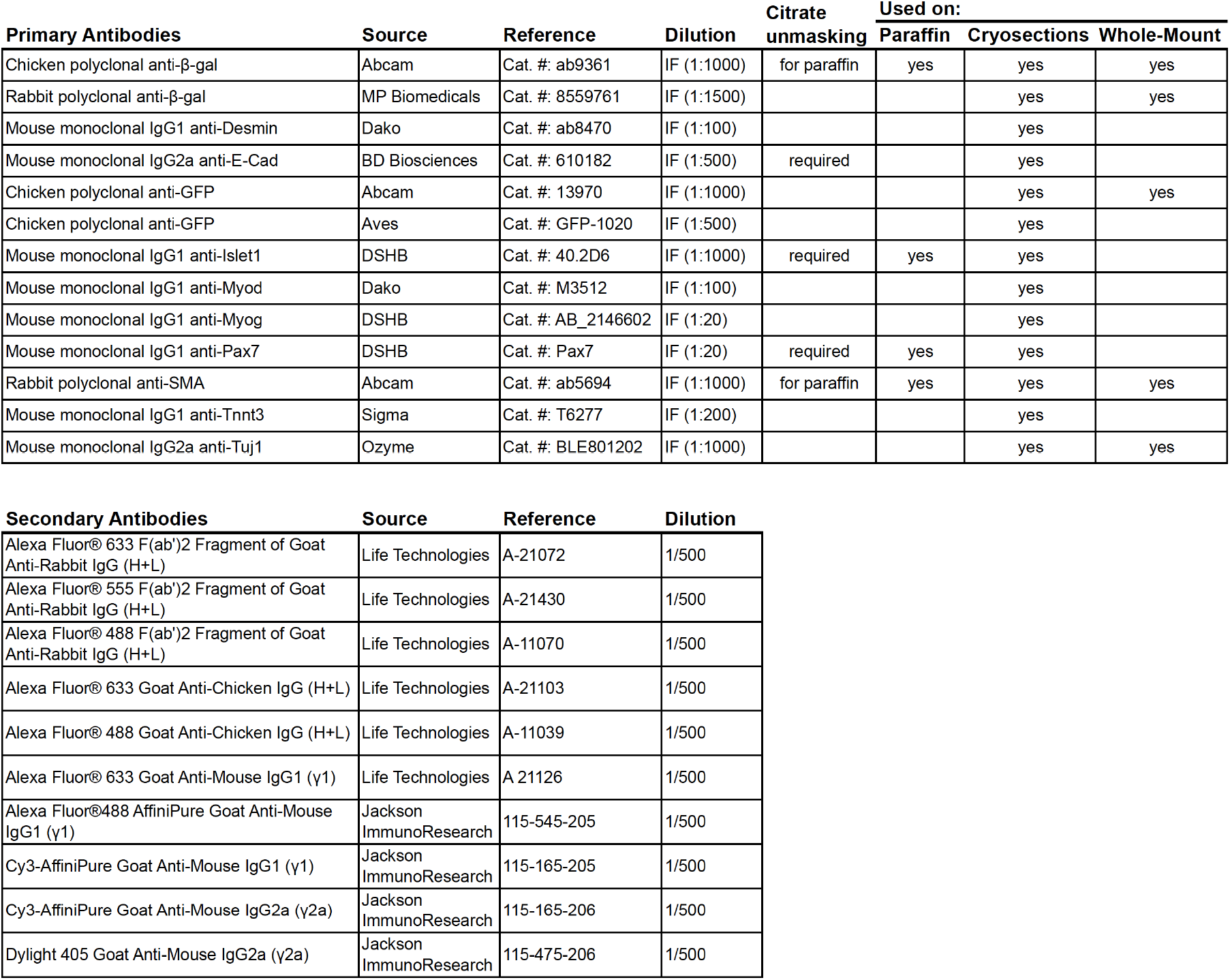
Antibodies used in this study

| Primary Antibodies | Source | Reference | Dilution | Citrate unmasking | Used on: |  |  |
| --- | --- | --- | --- | --- | --- | --- | --- |
|  |  |  |  |  | Paraffin | Cryosections | Whole-Mount |
| Chicken polyclonal anti-β-gal | Abcam | Cat. #: ab9361 | IF (1:1000) | for paraffin | yes | yes | yes |
| Rabbit polyclonal anti-β-gal | MP Biomedicals | Cat. #: 8559761 | IF (1:1500) | | | yes | yes |
| Mouse monoclonal IgG1 anti-Desmin | Dako | Cat. #: ab8470 | IF (1:100) |  |  | yes |  |
| Mouse monoclonal IgG2a anti-E-Cad | BD Biosciences | Cat. #: 610182 | IF (1:500) | required |  | yes |  |
| Chicken polyclonal anti-GFP | Abcam | Cat. #: 13970 | IF (1:1000) |  |  | yes | yes |
| Chicken polyclonal anti-GFP | Aves | Cat. #: GFP-1020 | IF (1:500) |  |  | yes |  |
| Mouse monoclonal IgG1 anti-Islet1 | DSHB | Cat. #: 40.2D6 | IF (1:1000) | required | yes | yes |  |
| Mouse monoclonal IgG1 anti-Myod | Dako | Cat. #: M3512 | IF (1:100) |  |  | yes |  |
| Mouse monoclonal IgG1 anti-Myog | DSHB | Cat. #: AB_2146602 | IF (1:20) |  |  | yes |  |
| Mouse monoclonal IgG1 anti-Pax7 | DSHB | Cat. #: Pax7 | IF (1:20) | required | yes | yes |  |
| Rabbit polyclonal anti-SMA | Abcam | Cat. #: ab5694 | IF (1:1000) | for paraffin | yes | yes | yes |
| Mouse monoclonal IgG1 anti-Tnnt3 | Sigma | Cat. #: T6277 | IF (1:200) |  |  | yes |  |
| Mouse monoclonal IgG2a anti-Tuj1 | Ozyme | Cat. #: BLE801202 | IF (1:1000) |  |  | yes | yes |

| Secondary Antibodies | Source | Reference | Dilution |
| --- | --- | --- | --- |
| Alexa Fluor® 633 F(ab')<sub>2</sub> Fragment of Goat Anti-Rabbit IgG (H+L) | Life Technologies | A-21072 | 1/500 |
| Alexa Fluor® 555 F(ab')<sub>2</sub> Fragment of Goat Anti-Rabbit IgG (H+L) | Life Technologies | A-21430 | 1/500 |
| Alexa Fluor® 488 F(ab')<sub>2</sub> Fragment of Goat Anti-Rabbit IgG (H+L) | Life Technologies | A-11070 | 1/500 |
| Alexa Fluor® 633 Goat Anti-Chicken IgG (H+L) | Life Technologies | A-21103 | 1/500 |
| Alexa Fluor® 488 Goat Anti-Chicken IgG (H+L) | Life Technologies | A-11039 | 1/500 |
| Alexa Fluor® 633 Goat Anti-Mouse IgG1 (γ1) | Life Technologies | A 21126 | 1/500 |
| Alexa Fluor®488 AffiniPure Goat Anti-Mouse IgG1 (γ1) | Jackson ImmunoResearch | 115-545-205 | 1/500 |
| Cy3-AffiniPure Goat Anti-Mouse IgG1 (γ1) | Jackson ImmunoResearch | 115-165-205 | 1/500 |
| Cy3-AffiniPure Goat Anti-Mouse IgG2a (γ2a) | Jackson ImmunoResearch | 115-165-206 | 1/500 |
| Dylight 405 Goat Anti-Mouse IgG2a (γ2a) | Jackson ImmunoResearch | 115-475-206 | 1/500 |

**Table2.**
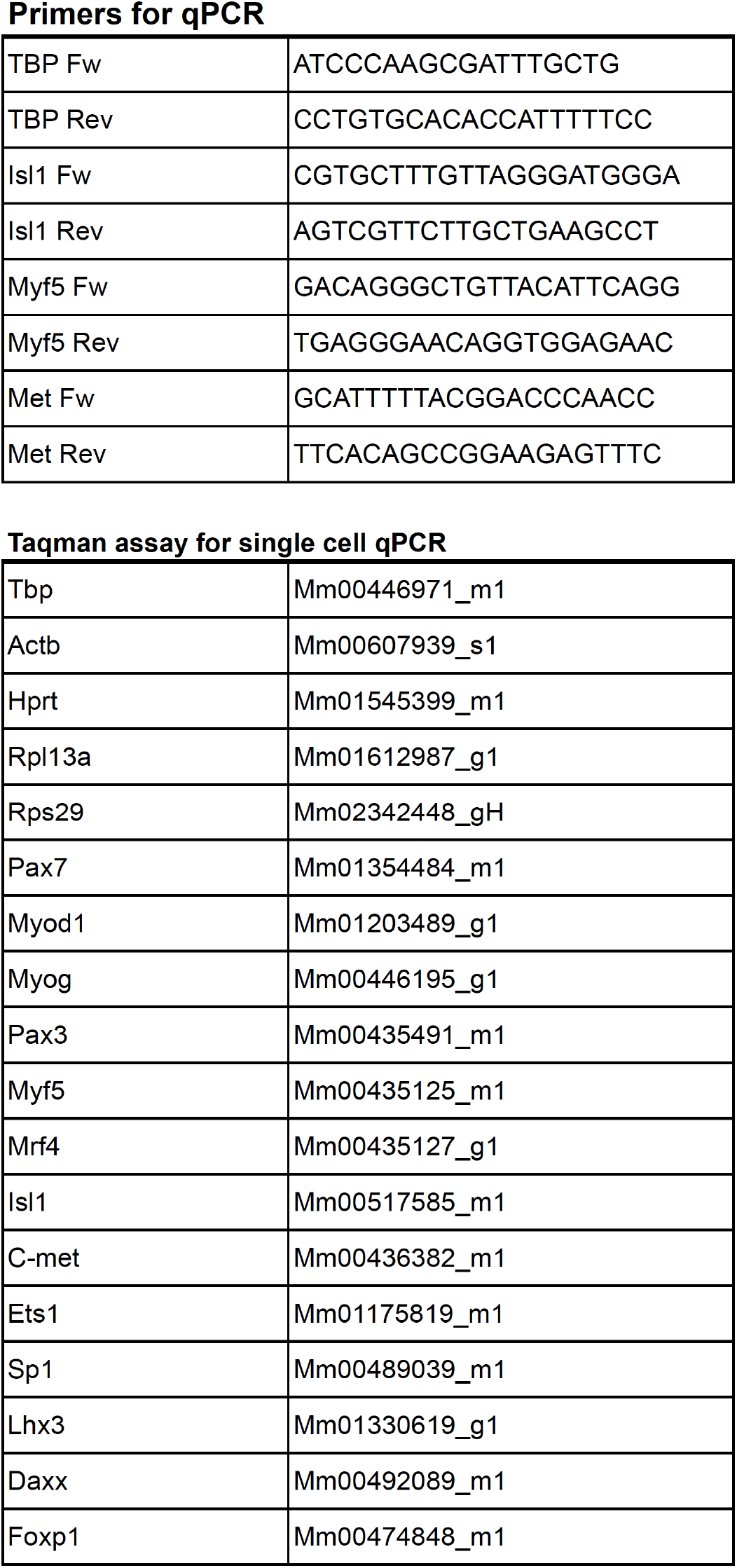
Primers used in this study

##### Convertion to relative expression

Raw Ct values were converted in relative expression using the following formula: Log2ex = LOD – Ct[Array] (Livak et al., 2013). With the LOD standing for the Limit Of Detection. When the Log2ex value obtained was negative (Ct[array]>LOD) the value was replaced by 0. To set up the LOD, we round the mean of the maximum Ct values for all the genes to the upper limit which gives a LOD of 21.

##### Normalization

The resulting relative expression values were normalized to the endogenous controls by substracting, for each cell, the average of its Actb, Rpl13, Rps29, and Hprt expression levels. An offset corresponding to the mean of all the calculated means was applied to all obtained values to avoid negative values.

##### Single cell filtering

From two independent experiments 66 cells were collected from the esophage. The criteria to keep a cell for further analysis were the following: i) to discard neurogenic progenitors, cells should not express Pax3 and/or Lhx3. ii) at least 4 out of the 5 positive control genes should be expressed, as well as at least 2 of the genes of interest. Applying this different filters 23 single cells were selected.

##### Correlation coefficient determination and p-value calculation

The Spearman’s rho correlation coefficient was calculated using the R function cor() with the ‘use’ parameter set at « pairwise.complete.obs », and all the null values previously replaced by NAs.

The coefficient correlation p-value was extracted from the cor.test() R function, using the same parameters.

### Data visualisation

The heatmap (Fig S5B) was generated using the pheatmap R package (pheatmap_1.0.10) with default parameters, and the correlogram (Fig 5F) was generated using the corrplot R package(corrplot_0.84), with the p-values mannually added. Violin plots in figure S1 were made in R using the ggplot2 package (ggplot2_3.1.0). R session info: R version 3.5.1 (2018-07-02), platform: x86_64-apple-darwin15.6.0 (64-bit), running under: macOS Sierra 10.12.6

### Static Imaging

Images were acquired using the following systems: Zeiss SteREO Discovery V20 microscope for whole embryos, a Zeiss Axioplan equipped with an Apotome and ZEN software (Carl Zeiss), Leica SPE or Leica TCS-SP8 with Leica Application Suite (LAS) software for tissue sections and a LSM 700 laser-scanning confocal microscope and ZEN software (Carl Zeiss) for tissue sections and whole mount immunostaining of cleared embryos. All images were assembled in Adobe Photoshop and InDesign (Adobe Systems). Volume-3D rendering on the z-stack series was performed in Imaris (version 7.2.1) software (Bitplane).

### Explant culture

Esophagi from E14.5 *Isl1*^*Cre*/+^*:R26*^*mTmG*/+^ embryos were micro-dissected leaving the stomach and pharyngeal muscles attached in RT HBSS (Gibco, 14025). The esophagi were immobilized on individual wells of 8 well glass bottom dishes (Ibidi, 80826) at the stomach end and pharyngeal ends using 0.3μl of Vetbond tissue adhesive (3M™, 1469SB). The explants were immediately embedded in a collagen matrix as previously reported (Placzek and Dale, 1999) with slight modifications. 700μl of collagenI (Corning, 354236), 200μl of reconstituted 5X DMEM-F12 (SIGMA, D2906) and 100μl neutralization buffer (50mM NaOH, 260mM NaHCO3, 200mM Hepes) were mixed throughrouly and kept on ice. 200μl of collagen matrix was added to each explant and allowed to polymerise for 10min in a culture incubator at 37°C, 5% CO2. Explant culture media was composed of Opti-MEM (Gibco, 51985-026) with 1%P/S and 20% FCS. 250ul of culture media containing Met inhibitors or the equivalent amount of DMSO (control) was added to each well and allowed to equilibrate for 30 min in a culture incubator. The Met inhibitors used were MGCD-265 (10μM, Selleck, 50mM stock in DMSO) and PF-0417903 (10-20μM, AbMole, 26.8mM stock in DMSO).

For static cultures, images of individual wells were acquired at 6 to 12h intervals on a Zeiss SteREO Discovery V20 microscope as Z-stacks and processed with the extended depth focus function on the Zen software.

For time-lapse imaging, the dish was placed in a microscope incubator chamber (37°C, 5% CO2) and the mGFP signal imaged with the 488 laser on an Leica TCS-SP8 inverted microscope, equipped with a HC PL APO CS2 10X/0.40 Dry objective and HyD hybrid detector (496-566nm). Confocal imaging of optical Z-planes (2.41μm) were acquired every 15 minutes over 14h using LAS X software. Z-stacks were projected as maximum intensity projection images, stitched and registered (linear registration) in Fiji. The migrating cells were tracked individually frame-by-frame using the ‘Manual Tracking’ plugin in Fiji. The following parameters were quantitated: total distance (μm, the distance covered by the whole track), velocity (μm/min, ratio between the total distance and total time of the track), displacement (μm, the lenght of the resultant vector between ti and tf of the track), efficiency (ratio between displacement and total distance), net velocity (μm/min, ratio between the displacement and total time of the track).

### Quantitation of muscle area

The muscle area on transverse esophagus cryosections (Suppl. Figure 1I) was quantified on Fiji. Channels were split threshold levels adjusted on the Tnnt3 channel. The freehand selection tool was used to trace the outline of each esophagus crossection (referred as Region of interest, ROI). Threshold levels were kept constant for all samples. The Analyze/Measure tool was set to calculate the area of the ROI limited to the threshold for the Tnnt3 channel.

### EdU Administration In Vivo

For proliferation experiments in vivo, 5-ethyl-20-deoxyuridine (EdU; Invitrogen E10187) was injected intraperitoneally and detected as described in (Comai et al., 2014).

## ACKNOWLEDGEMENTS

We acknowledge funding support from the Institut Pasteur, Association Française contre le Myopathies, Agence Nationale de la Recherche (Laboratoire d’Excellence Revive, Investissement d’Avenir; ANR-10-LABX-73). We acknowledge the service of Pasteur Imaging platform (PBI), Pasteur Mouse Genetic Engineering platform (CIGM) and Pasteur Flow Cytometry Platform (CEITEC). We also thank C. Cimper for technical assistance.

## AUTHOR CONTRIBUTIONS

G.C., E.H., S.G. and S.T. conceived and designed the experiments and wrote the manuscript. G.C., E.H. and S.G. performed *Met^D^* and *Hgf* mutant analysis. E.H and F.P. performed single cell RT-PCR analysis. S.M. performed bioinformatic analysis. E.H. and G.C. performed chimera analysis. S.P. and E.H performed ES derivation. G.C. performed ISH (Rnascope®) and whole mount immunostainings. M.G. and G.K. contributed to *Hgf* mutant analysis. All authors interpreted the results, read and approved the final manuscript.

---

**Supplementary Figure 1.**
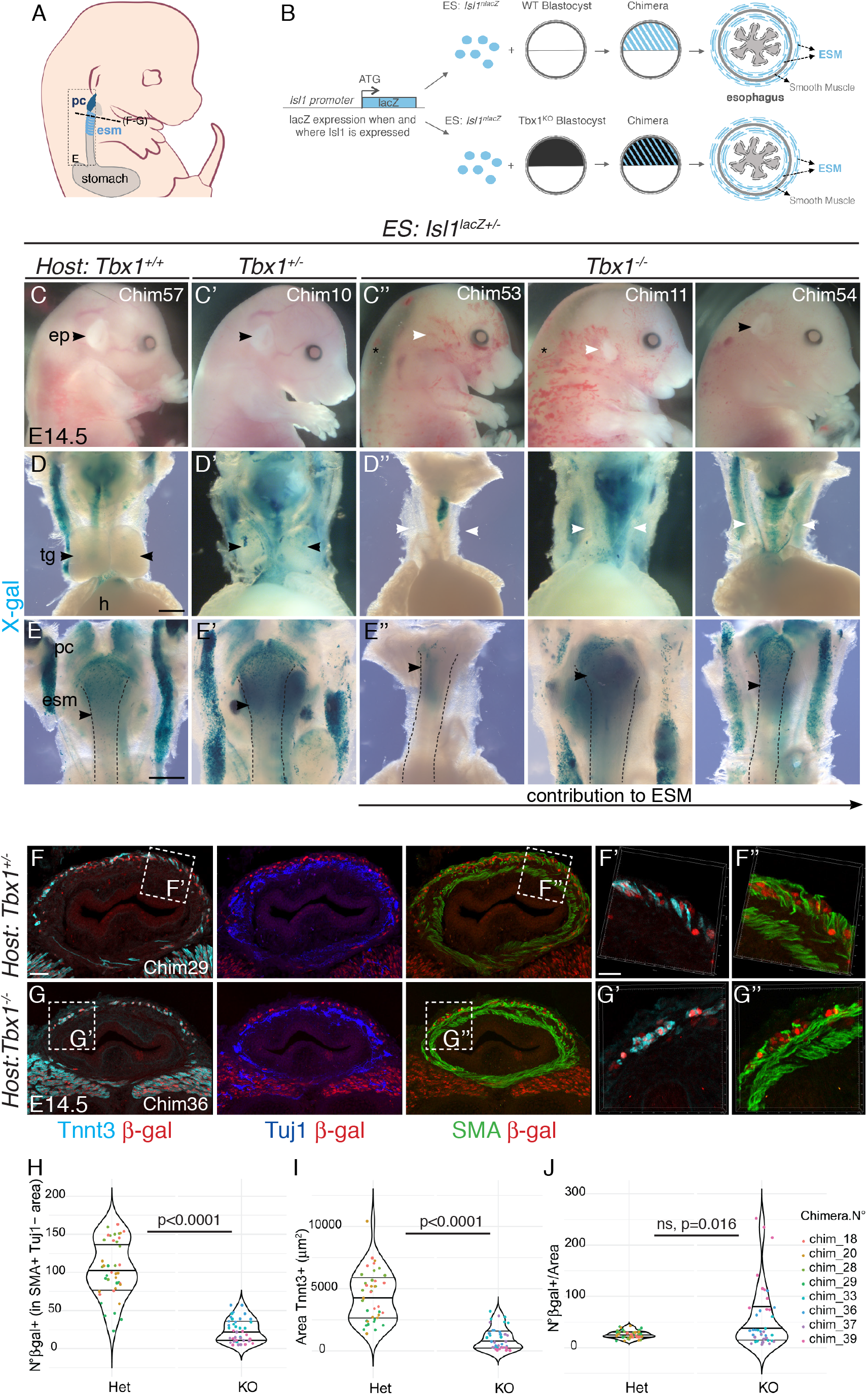
*Isl1*^*nlacZ*/+^ cells colonize the esophagus of chimeric *Tbx1* mutants. (A) Structures and levels analysed in the study. (B) Schematic summary of the chimera experiment. (C-C’’) Macroscopic views of E14.5/E15.5 *Isl1*^*nlacZ*/+^ ES cell derived chimeric embryos (C, *Tbx1* wildtype host; C’, *Tbx1* heterozygote mutant host; C’’, *Tbx1* homozygote knockout host). A black arrowhead indicates a normal and a white arrowhead a reduced or absent outer ear pinna. Chimeras of *Tbx1*^-/-^ host are generally edemic (C’’, asterisk). (D-E) X-gal staining of dissected digestive tracks including the heart and thymus glands. A black arrowhead indicates normal (D, D’) and a white arrowhead absent thymus glands (D’’). (E-E’’) Dorsal views of the esophagus (dotted lines). *Isl1*^*nlacZ*/+^ ES contribute extensively to wild type (E) and heterozygote (CE’) chimeric embryos, while they contribute to different extent in *Tbx1*^-/-^ hosts (E’’). A black arrowhead in E-E’’ indicates the most caudal level to which *Isl1*^*nlacZ*/+^ ES contribute. (F-G) Immunostainings on transverse cryosections at the esophagus level of E14.5/E15.4 *Isl1*^*nlacZ*/+^ ES cell derived chimeric embryos for Tuj1 (neurons), SMA (smooth muscle actin) and β-gal (Isl1-expressing cells). β-gal+ cells are present in the smooth muscle scaffold and contribute to Tnnt3+ fibers in both control (F’, F”) and *Tbx1*^-/-^ hosts (G’, G”). (H-J) Quantitative analysis (violin plots) of the number of β-gal+ cells per section in present in the smooth muscle scaffold (H), the Tnnt3+ Area per section (I) and ratio of β-gal+ cells per muscle are per section (J) in chimeras derived from heterozygote and Tbx1 mutant hosts. Dots, quantification on individual sections. p= p-value of Mann-Whitney test. Center lines show the medians; box limits indicate the 25th and 75th percentiles as determined by R software. ep, ear pinna; esm, esophagus striated muscle; h, heart; pc, pharyngeal constrictor; tg, thymus glands. Scale bars: B,C, 500μm; D, 40μm, D’, 20μm Scale bars: D,E, 500μm; F, 40μm, F’, 20μm

**Supplementary Figure 2.**
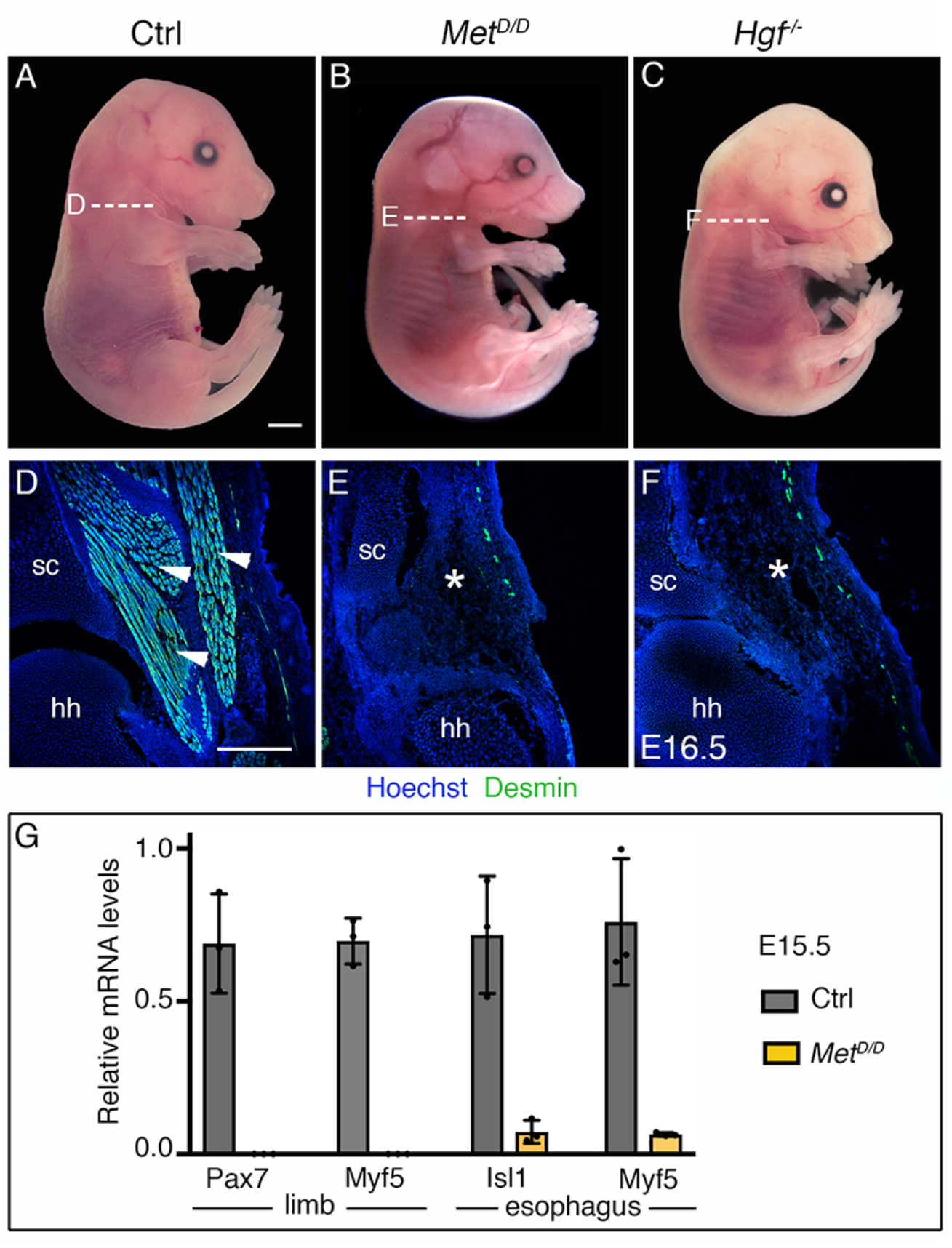
Phenotype of *Met* and *Hgf* mutants. (A-C) Macroscopic view of E16.5 control, *Met^D/D^* and *Hgf*^-/-^ foetuses (D-F) Immunostainings on coronal cryosections at levels indicated in (A-C) for the myogenic markers Desmin. Note the absence of limb musculature in *Met* and *Hgf* mutants (white asterisks) (n=3 each conditions). (G) RT-qPCR analysis for *Pax7* and *Myf5* in limbs and for *Isl1* and *Myf5* in esophagi of E15.5 control and *Met^D/D^* foetuses. All data points are plotted and presented as the mean ± SEM (error bars) (n=3 each condition). Abreviations: hh, humeral head; sc, scapula. Scale bars: A, 1000 μm; D, 200 μm.

**Supplementary Figure 3.**
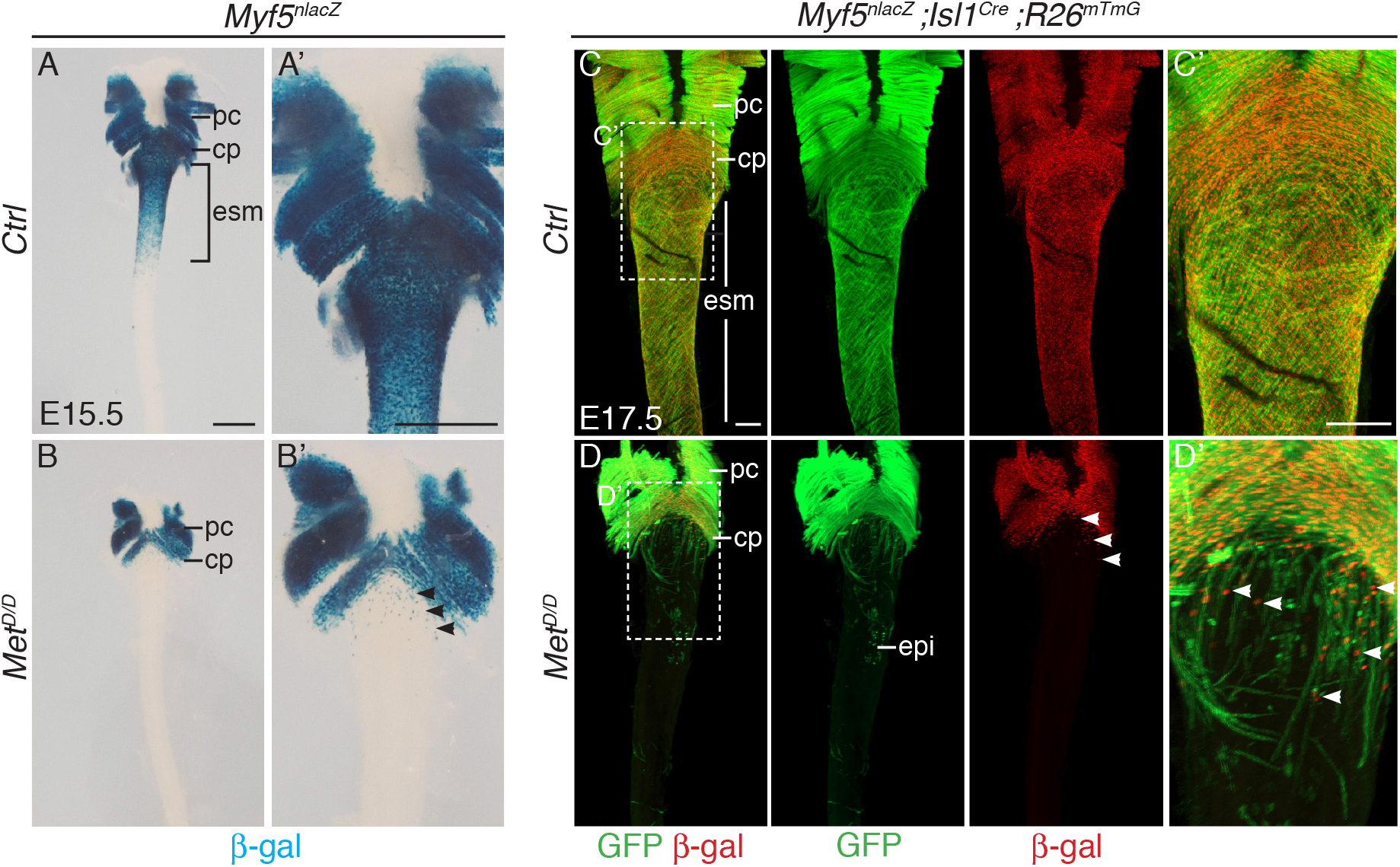
Myogenic cells are present anteriorly in the esophagus of *Met^D/D^* mutants. (A-B) Whole mount X-gal staining of the upper esophagus of E15.5 Met mutant and control embryos (dorsal view). (A’-B’). Higher magnification views of the images in (A-B). Black arrowheads point to *Myf5*^*nlacZ*/+^ progenitor cells present in the upper esophagus in the mutant. C-D. Whole mount immunostaining of the upper esophagus of E17.5 Met mutant and control embryos (dorsal views) stained for β-gal (Myf5 progenitors) and GFP (Isl1 lineage tracing). *Myf5*^*nlacZ*/+^ progenitor cells are present in the anteriormost esophagus of the mutant (D, D’, white arrowheads). cp, cricopharyngeous; epi, epithelial *Isl1*-derived cells; esm, esophagus striated muscle; pc, pharyngeal constrictor. Scale bars: A, A’, 500μm; C, C’ 200μm

**Supplementary Figure 4.**
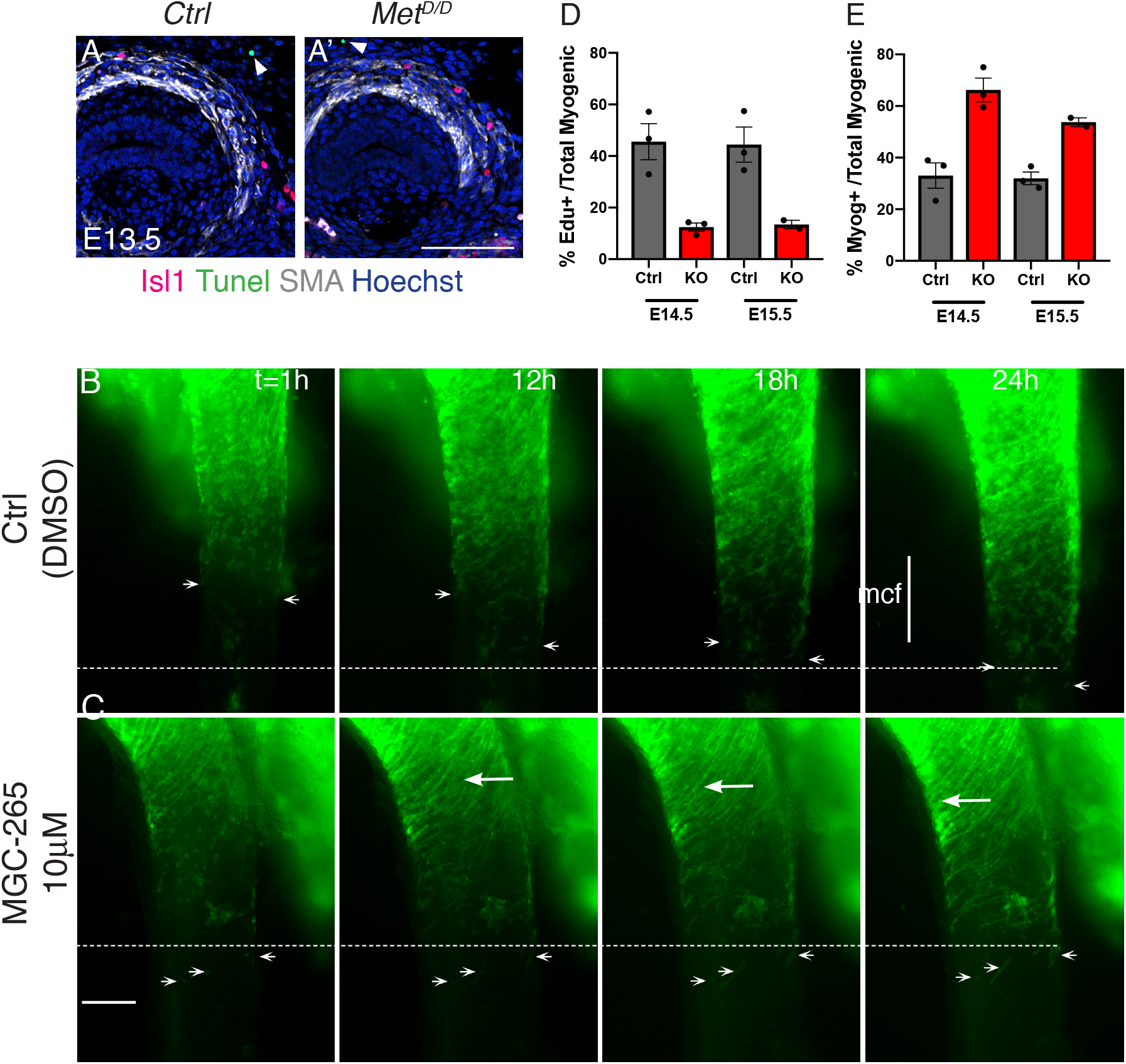
*Met* invalidation does not affect the proliferation and survival of *Isl1* progenitors. (A-A’) Tunnel assay on transverse cryosections of E15.5 Met mutant and control embryos. White arrowheads point to Tunnel+ cell outside the esm. (B-C) Time series from a static esophagus explant culture experiment in the presence of DMSO (ctrl, A) or 10μM MGCD-265 (B). White arrowheads point to *Isl1*-derived progenitor cells present at the mononucleated cell front (mcf). White arrows highlight the high numbers of fibers that appear progresively in the inhibitor condition. Time (t) is indicated in hours. Dotted lines show the location of the mcf at 24h in the control and inhibitor condition. (D-E) Quantification of the % myogenic cells that are Edu+ (proliferative) and Myog+ (ongoing differentiation) (n=3). Scale bars: A’, 50 μm; B, 200μm

**Supplementary Figure 5.**
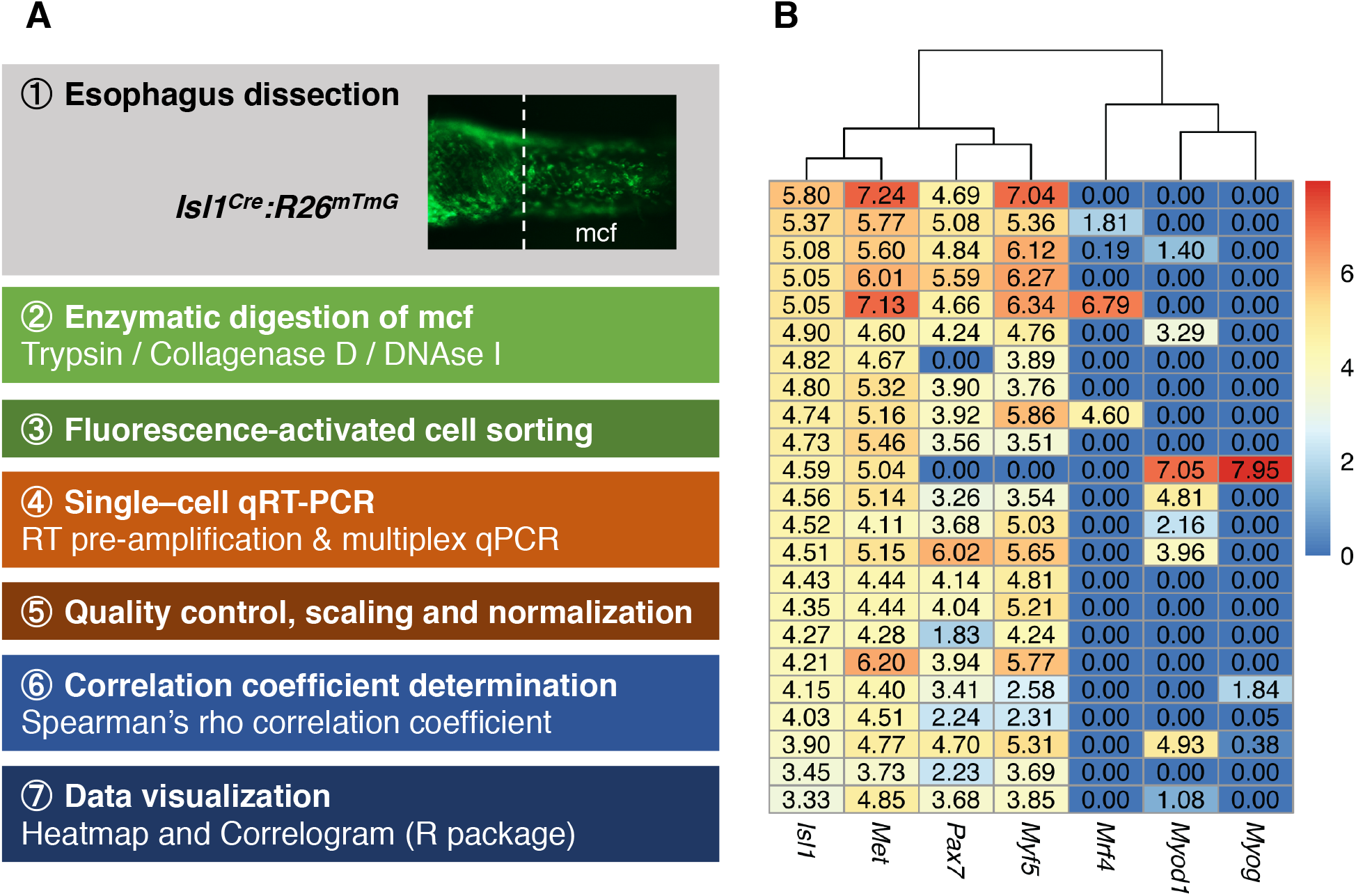
Esophagus single cell analysis. (A) Experimental procedure for the esophagus mononucleated cells analysis. The heatmap shows the normalized converted expression values, with columns corresponding to the genes and rows corresponding to the different single cells. Dendrogram shows the hierarchical clustering using the euclidean distance between genes.

## RICH MEDIA FILES

**Movie S1**. Time lapse movie of a control E14.5 *Isl1^Cre^;R26^mTmG^* esophagus explant culture. Time as hh:mm. Related to Figure4. Individual GFP+ cells are tracked on different colors.

**Movie S2**. Time lapse movie of a E14.5 *Isl1^Cre^;R26^mTmG^* esophagus explant culture treated with 10μM PF-0417903. Time as hh:mm. Related to Figure4. Individual GFP+ cells are tracked on different colors.

**Movie S3**. Time lapse movie of a E14.5 *Isl1^Cre^;R26^mTmG^* esophagus explant culture treated with 10μM MGCD-265. Time as hh:mm. Related to Figure4. Individual GFP+ cells are tracked on different colors.

## REFERENCES

Arnold, J.S., Werling, U., Braunstein, E.M., Liao, J., Nowotschin, S., Edelmann, W., Hebert, J.M., and Morrow, B.E. (2006). Inactivation of Tbx1 in the pharyngeal endoderm results in 22q11DS malformations. Development 133, 977–987.

Birchmeier, C., and Gherardi, E. (1998). Developmental roles of HGF/SF and its receptor, the c-Met tyrosine kinase. Trends in cell biology 8, 404–410.

Biressi, S., Molinaro, M., and Cossu, G. (2007). Cellular heterogeneity during vertebrate skeletal muscle development. Developmental biology 308, 281–293.

Bladt, F., Riethmacher, D., Isenmann, S., Aguzzi, A., and Birchmeier, C. (1995). Essential role for the c-met receptor in the migration of myogenic precursor cells into the limb bud. Nature 376, 768–771.

Bonnet, A., Dai, F., Brand-Saberi, B., and Duprez, D. (2010). Vestigial-like 2 acts downstream of MyoD activation and is associated with skeletal muscle differentiation in chick myogenesis. Mechanisms of development 127, 120–136.

Brand-Saberi, B., Muller, T.S., Wilting, J., Christ, B., and Birchmeier, C. (1996). Scatter factor/hepatocyte growth factor (SF/HGF) induces emigration of myogenic cells at interlimb level in vivo. Developmental biology 179, 303–308.

Cai, C.L., Liang, X., Shi, Y., Chu, P.H., Pfaff, S.L., Chen, J., and Evans, S. (2003). Isl1 identifies a cardiac progenitor population that proliferates prior to differentiation and contributes a majority of cells to the heart. Developmental cell 5, 877–889.

Chihara, D., Romer, A.I., Bentzinger, C.F., Rudnicki, M.A., and Krauss, R.S. (2015). PAX7 is required for patterning the esophageal musculature. Skeletal muscle 5, 39.

Comai, G., Sambasivan, R., Gopalakrishnan, S., and Tajbakhsh, S. (2014). Variations in the efficiency of lineage marking and ablation confound distinctions between myogenic cell populations. Developmental cell 31, 654–667.

Comai, G., and Tajbakhsh, S. (2014). Molecular and cellular regulation of skeletal myogenesis. Current topics in developmental biology 110, 1–73.

Das, P., and May, C.L. (2011). Expression analysis of the Islet-1 gene in the developing and adult gastrointestinal tract. Gene expression patterns: GEP 11, 244–254.

Dietrich, S., Abou-Rebyeh, F., Brohmann, H., Bladt, F., Sonnenberg-Riethmacher, E., Yamaai, T., Lumsden, A., Brand-Saberi, B., and Birchmeier, C. (1999). The role of SF/HGF and c-Met in the development of skeletal muscle. Development 126, 1621–1629.

Diogo, R., Kelly, R.G., Christiaen, L., Levine, M., Ziermann, J.M., Molnar, J.L., Noden, D.M., and Tzahor, E. (2015). A new heart for a new head in vertebrate cardiopharyngeal evolution. Nature 520, 466–473.

Emery, A.E. (2002). The muscular dystrophies. Lancet 359, 687–695.

Epstein, J.A., Shapiro, D.N., Cheng, J., Lam, P.Y., and Maas, R.L. (1996). Pax3 modulates expression of the c-Met receptor during limb muscle development. Proceedings of the National Academy of Sciences of the United States of America 93, 4213–4218.

Gopalakrishnan, S., Comai, G., Sambasivan, R., Francou, A., Kelly, R.G., and Tajbakhsh, S. (2015). A Cranial Mesoderm Origin for Esophagus Striated Muscles. Developmental cell 34, 694–704.

Harel, I., Nathan, E., Tirosh-Finkel, L., Zigdon, H., Guimaraes-Camboa, N., Evans, S.M., and Tzahor, E. (2009). Distinct origins and genetic programs of head muscle satellite cells. Developmental cell 16, 822–832.

Heude, E., Tesarova, M., Sefton, E.M., Jullian, E., Adachi, N., Grimaldi, A., Zikmund, T., Kaiser, J., Kardon, G., Kelly, R.G., et al. (2018). Unique morphogenetic signatures define mammalian neck muscles and associated connective tissues. eLife 7.

Jerome, L.A., and Papaioannou, V.E. (2001). DiGeorge syndrome phenotype in mice mutant for the T-box gene, Tbx1. Nature genetics 27, 286–291.

Kassar-Duchossoy, L., Giacone, E., Gayraud-Morel, B., Jory, A., Gomes, D., and Tajbakhsh, S. (2005). Pax3/Pax7 mark a novel population of primitive myogenic cells during development. Genes & development 19, 1426–1431.

Kelly, R.G., Jerome-Majewska, L.A., and Papaioannou, V.E. (2004). The del22q11.2 candidate gene Tbx1 regulates branchiomeric myogenesis. Human molecular genetics 13, 2829–2840.

Krauss, R.S., Chihara, D., and Romer, A.I. (2016). Embracing change: striated-for-smooth muscle replacement in esophagus development. Skeletal muscle 6, 27.

Lescroart, F., Hamou, W., Francou, A., Theveniau-Ruissy, M., Kelly, R.G., and Buckingham, M. (2015). Clonal analysis reveals a common origin between nonsomite-derived neck muscles and heart myocardium. Proceedings of the National Academy of Sciences of the United States of America 112, 1446–1451.

Livak, K.J., Wills, Q.F., Tipping, A.J., Datta, K., Mittal, R., Goldson, A.J., Sexton, D.W., and Holmes, C.C. (2013). Methods for qPCR gene expression profiling applied to 1440 lymphoblastoid single cells. Methods 59, 71–79.

Maina, F., Casagranda, F., Audero, E., Simeone, A., Comoglio, P.M., Klein, R., and Ponzetto, C. (1996). Uncoupling of Grb2 from the Met receptor in vivo reveals complex roles in muscle development. Cell 87, 531–542.

Maina, F., Pante, G., Helmbacher, F., Andres, R., Porthin, A., Davies, A.M., Ponzetto, C., and Klein, R. (2001). Coupling Met to specific pathways results in distinct developmental outcomes. Molecular cell 7, 1293–1306.

Muzumdar, M.D., Tasic, B., Miyamichi, K., Li, L., and Luo, L. (2007). A global double-fluorescent Cre reporter mouse. Genesis 45, 593–605.

Nathan, E., Monovich, A., Tirosh-Finkel, L., Harrelson, Z., Rousso, T., Rinon, A., Harel, I., Evans, S.M., and Tzahor, E. (2008). The contribution of Islet1-expressing splanchnic mesoderm cells to distinct branchiomeric muscles reveals significant heterogeneity in head muscle development. Development 135, 647–657.

Pfaff, S.L., Mendelsohn, M., Stewart, C.L., Edlund, T., and Jessell, T.M. (1996). Requirement for LIM homeobox gene Isl1 in motor neuron generation reveals a motor neuron-dependent step in interneuron differentiation. Cell 84, 309–320.

Placzek, M., and Dale, K. (1999). Tissue recombinations in collagen gels. Methods Mol Biol 97, 293–304.

Prunotto, C., Crepaldi, T., Forni, P.E., Ieraci, A., Kelly, R.G., Tajbakhsh, S., Buckingham, M., and Ponzetto, C. (2004). Analysis of Mlc-lacZ Met mutants highlights the essential function of Met for migratory precursors of hypaxial muscles and reveals a role for Met in the development of hyoid arch-derived facial muscles. Developmental dynamics: an official publication of the American Association of Anatomists 231, 582–591.

Randolph, M.E., and Pavlath, G.K. (2015). A muscle stem cell for every muscle: variability of satellite cell biology among different muscle groups. Frontiers in aging neuroscience 7, 190.

Relaix, F., Montarras, D., Zaffran, S., Gayraud-Morel, B., Rocancourt, D., Tajbakhsh, S., Mansouri, A., Cumano, A., and Buckingham, M. (2006). Pax3 and Pax7 have distinct and overlapping functions in adult muscle progenitor cells. The Journal of cell biology 172, 91–102.

Relaix, F., Rocancourt, D., Mansouri, A., and Buckingham, M. (2005). A Pax3/Pax7-dependent population of skeletal muscle progenitor cells. Nature 435, 948–953.

Rishniw, M., Xin, H.B., Deng, K.Y., and Kotlikoff, M.I. (2003). Skeletal myogenesis in the mouse esophagus does not occur through transdifferentiation. Genesis 36, 81–82.

Romer, A.I., Singh, J., Rattan, S., and Krauss, R.S. (2013). Smooth muscle fascicular reorientation is required for esophageal morphogenesis and dependent on Cdo. The Journal of cell biology 201, 309–323.

Rudnicki, M.A., Schnegelsberg, P.N., Stead, R.H., Braun, T., Arnold, H.H., and Jaenisch, R. (1993). MyoD or Myf-5 is required for the formation of skeletal muscle. Cell 75, 1351–1359.

Sambasivan, R., Gayraud-Morel, B., Dumas, G., Cimper, C., Paisant, S., Kelly, R.G., and Tajbakhsh, S. (2009). Distinct regulatory cascades govern extraocular and pharyngeal arch muscle progenitor cell fates. Developmental cell 16, 810–821.

Sambasivan, R., Kuratani, S., and Tajbakhsh, S. (2011). An eye on the head: the development and evolution of craniofacial muscles. Development 138, 2401–2415.

Scaal, M., Bonafede, A., Dathe, V., Sachs, M., Cann, G., Christ, B., and Brand-Saberi, B. (1999). SF/HGF is a mediator between limb patterning and muscle development. Development 126, 4885–4893.

Schmidt, C., Bladt, F., Goedecke, S., Brinkmann, V., Zschiesche, W., Sharpe, M., Gherardi, E., and Birchmeier, C. (1995). Scatter factor/hepatocyte growth factor is essential for liver development. Nature 373, 699–702.

Schmittgen, T.D., Lee, E.J., Jiang, J., Sarkar, A., Yang, L., Elton, T.S., and Chen, C. (2008). Real-time PCR quantification of precursor and mature microRNA. Methods 44, 31–38.

Schubert, F.R., Singh, A.J., Afoyalan, O., Kioussi, C., and Dietrich, S. (2018). To roll the eyes and snap a bite - function, development and evolution of craniofacial muscles. Seminars in cell & developmental biology.

Sheehan, N.J. (2008). Dysphagia and other manifestations of oesophageal involvement in the musculoskeletal diseases. Rheumatology (Oxford) 47, 746–752.

Srinivas, S., Watanabe, T., Lin, C.S., William, C.M., Tanabe, Y., Jessell, T.M., and Costantini, F. (2001). Cre reporter strains produced by targeted insertion of EYFP and ECFP into the ROSA26 locus. BMC developmental biology 1, 4.

Sun, Y., Liang, X., Najafi, N., Cass, M., Lin, L., Cai, C.L., Chen, J., and Evans, S.M. (2007). Islet 1 is expressed in distinct cardiovascular lineages, including pacemaker and coronary vascular cells. Developmental biology 304, 286–296.

Tabler, J.M., Rigney, M.M., Berman, G.J., Gopalakrishnan, S., Heude, E., Al-Lami, H.A., Yannakoudakis, B.Z., Fitch, R.D., Carter, C., Vokes, S., et al. (2017). Cilia-mediated Hedgehog signaling controls form and function in the mammalian larynx. eLife 6.

Tajbakhsh, S. (2009). Skeletal muscle stem cells in developmental versus regenerative myogenesis. J Intern Med 266, 372–389.

Tajbakhsh, S., Rocancourt, D., and Buckingham, M. (1996). Muscle progenitor cells failing to respond to positional cues adopt non-myogenic fates in myf-5 null mice. Nature 384, 266–270.

Tajbakhsh, S., Rocancourt, D., Cossu, G., and Buckingham, M. (1997). Redefining the genetic hierarchies controlling skeletal myogenesis: Pax-3 and Myf-5 act upstream of MyoD. Cell 89, 127–138.

Tam, P.P., and Rossant, J. (2003). Mouse embryonic chimeras: tools for studying mammalian development. Development 130, 6155–6163.

Trusolino, L., Bertotti, A., and Comoglio, P.M. (2010). MET signalling: principles and functions in development, organ regeneration and cancer. Nature reviews Molecular cell biology 11, 834–848.

Wang, F., Flanagan, J., Su, N., Wang, L.C., Bui, S., Nielson, A., Wu, X., Vo, H.T., Ma, X.J., and Luo, Y. (2012). RNAscope: a novel in situ RNA analysis platform for formalin-fixed, paraffin-embedded tissues. The Journal of molecular diagnostics: JMD 14, 22–29.

Webster, M.T., and Fan, C.M. (2013). c-MET regulates myoblast motility and myocyte fusion during adult skeletal muscle regeneration. PloS one 8, e81757.

Yang, X.M., Vogan, K., Gros, P., and Park, M. (1996). Expression of the met receptor tyrosine kinase in muscle progenitor cells in somites and limbs is absent in Splotch mice. Development 122, 2163–2171.

Yokomizo, T., Yamada-Inagawa, T., Yzaguirre, A.D., Chen, M.J., Speck, N.A., and Dzierzak, E. (2012). Whole-mount three-dimensional imaging of internally localized immunostained cells within mouse embryos. Nature protocols 7, 421–431.

Zhang, Z., Huynh, T., and Baldini, A. (2006). Mesodermal expression of Tbx1 is necessary and sufficient for pharyngeal arch and cardiac outflow tract development. Development 133, 3587–3595.

Zhao, W., and Dhoot, G.K. (2000). Both smooth and skeletal muscle precursors are present in foetal mouse oesophagus and they follow different differentiation pathways. Developmental dynamics: an official publication of the American Association of Anatomists 2l8, 587–602.

